# RICTOR regulates an interspecies crosstalk that influences longevity through a novel methionine cycle-mitophagy axis

**DOI:** 10.1101/2025.07.11.664440

**Authors:** Simran Motwani, Somya Bhandari, Shivani Chitkara, Rajat Ujjainiya, Shantanu Sengupta, Arnab Mukhopadhyay

## Abstract

Adaptive modulation of physiological traits in response to environmental variability, particularly dietary fluctuations, is essential for organismal fitness. Such adaptability is governed by complex gene-diet interactions, yet the molecular circuits integrating microbe-derived metabolites with host metabolic and stress response pathways remain less explored. Here, we identify the conserved mechanistic target of rapamycin complex 2 (mTORC2) component, RICTOR, as a critical regulator of dietary plasticity in *Caenorhabditis elegans*, specifically in response to bacterially derived vitamin B12 (B12). Loss of *rict-1*, the *C. elegans* ortholog of RICTOR, confers enhanced osmotic stress tolerance and longevity on B12-rich bacterial diets. These phenotypic adaptations require two B12-dependent enzymes: methionine synthase (METR-1), functioning in the folate-methionine cycle (Met-C), and methylmalonyl-CoA mutase (MMCM-1), a mitochondrial enzyme essential for propionate catabolism. The latter catalyzes the formation of succinyl-CoA, subsequently converted to succinate via the tricarboxylic acid (TCA) cycle. Elevated succinate levels were found to induce mitochondrial fragmentation, thereby activating mitophagy, an autophagic process indispensable for the increased stress resilience and longevity observed in the *rict-1* mutants. Crucially, this Met-C-mitophagy axis is modulated by microbial inputs, with B12 and methionine acting as proximal dietary signals. Our findings delineate a mechanistic framework through which RICTOR restrains host sensitivity to microbial-derived metabolites, thus maintaining mitochondrial homeostasis and regulating lifespan. This work reveals a pivotal role for RICTOR in insulating host physiology from environmental nutrient-driven perturbations by modulating organellar quality control pathways.

## Introduction

Diet has a profound effect on the life history traits of an organism, including longevity. Organisms possess intrinsic homeostatic mechanisms that maintain a normal lifespan despite exposure to diverse nutritional inputs; this is important as abrupt changes in life history traits on fluctuating food availability may have serious consequences on species survival [1]. This adaptive capacity to diet of different nutritional values is maintained by gene-diet interactions, where host genetics counters dietary inputs to maintain normal life history traits [2]. However, in some genetic mutants studied in laboratories, variations in lifespan or other phenotypes become apparent only on specific diets, thereby giving us a window of opportunity to understand how diet-gene interactions maintain normal lifespan and how their perturbations may lead to diseases.

The bacterivorous nematode, *C. elegans*, has been instrumental in investigating molecular mechanisms of some of these gene-diet pairs [3]. The wild-type worms are resistant to changes in the quality of bacterial diet that they are fed on in the laboratory, and do not show dramatic changes in life history traits. But often genetic mutants are serendipitously found to exhibit altered phenotypes only on a specific diet. For example, when the *alh-6* (a proline catabolism gene) mutant is fed the high proline-containing *E. coli* OP50 diet, it cannot convert the proline-5-carboxylate (P5C) to glutamate and fails to maintain mitochondrial homeostasis. As a result, the mutants generate ROS and have damaged mitochondria on OP50, but not on the HT115 diet, leading to accelerated aging [4]. The nuclear hormone receptor gene *nhr-114*, when mutated, shows sterility only on OP50 but not on HT115, as the latter is enriched in the essential amino acid tryptophan, which aids in protecting germline through a possible detoxification mechanism [5]. Further, the neuromedin U receptor gene *nmur-1* mutant exhibits food-type dependent lifespan by sensing the difference in the lipopolysaccharide structure between bacteria, thus lives longer on OP50 as compared to HT115 [6]. Our lab has identified a novel gene-diet pair where the kinase-dead mutant, *flr-4(n2259)* has enhanced flux through the folate and methionine cycle (Met-C) leading to the activation of the p38-MAPK pathway and, in turn, differential expression of xenobiotic genes in response to elevated B12 levels in HT115, leading to increased lifespan [7, 8]. Similarly, a mitochondrial ribosomal gene, *mrpl-2* mutant showed B12-dependent lifespan and healthspan benefits by regulating the mitochondrial unfolded protein response [9]. When the worm ortholog of the mTORC2 subunit RICTOR gene, *rict-1*, is mutated, it leads to increased lifespan on HT115 but not on OP50 diet, dependent on the activation of the transcription factor NRF2/SKN-1 [10, 11]. Interestingly, although RICTOR/mTORC2 is an important kinase complex that lies at the crossroads of numerous signaling pathways, the detailed mechanism of how it is involved in adaptive capacity to different diets is less known.

In this study, we show that the *C. elegans* RICT-1 is a part of a gene-diet interaction that maintains adaptive capacity to diets with different B12 content. The mTORC2 complex is a conserved regulator that directs a wide range of fundamental processes such as cell survival, proliferation, and metabolism by integrating environmental and intracellular cues. Depletion of RICTOR causes the disruption of mTORC2 assembly and activity, implying that it plays a critical role in the integrity and stabilization of mTORC2 [12]. Much emerging evidence demonstrates that mTORC2 plays an essential role in glucose metabolism and homeostasis [13–15]. mTORC2-activated glucose metabolism can supply critical resources for nucleotide biosynthesis [16]. Aside from being triggered by ROS, mTORC2 is also engaged in ROS metabolism via regulating glutathione and NADPH production [15]. mTORC2 also plays a crucial role in lipid metabolism, boosting lipogenesis while decreasing lipolysis and fatty acid oxidation [17, 18]. Considering the central role of mTORC2, its dysregulation is linked to human disorders such as type 2 diabetes mellitus and cancer [19, 20].

Here, we show that the *rict-1* mutant grown on a high B12 diet engages Met-C as well as the transsulfuration and the canonical propionate catabolic pathways. This leads to an increase in the levels of succinate that fragments the mitochondria to signal the increase in the prolongevity process of mitophagy. Interestingly, we show that the bacterial diet supplies both B12 as well as the essential amino acid methionine (Met) to augment succinate production, mitochondrial fragmentation, mitophagy, and promote stress tolerance and longevity. Together, our study elucidates an elegant mechanism of inter-species communication that regulates stress tolerance and longevity of the host, dependent on nutrients supplied by the microbiota, an axis modulated by RICTOR to prevent abrupt changes in lifespan.

## Results

### Vitamin B12-driven stress tolerance and lifespan benefits in the *rict-1* mutant

The dietary makeup of bacteria plays a significant role in shaping the longevity and developmental traits of *rict-1* loss-of-function mutants, such as slowed growth and reduced body size [10, 11]. As reported previously, we also found that the *rict-1(ft7)* [*rict-1(-)*] worms display an extended lifespan when grown on *E. coli* HT115, compared to *E. coli* OP50 (**Figure 1A**). Additionally, we found that *rict-1(-)* worms fed with the HT115 diet exhibit heightened resistance to osmotic stress and improved recovery kinetics compared to the wild-type (**Figure 1B**). The wild-type worms remain unaffected by these dietary alterations.

**Figure 1:**
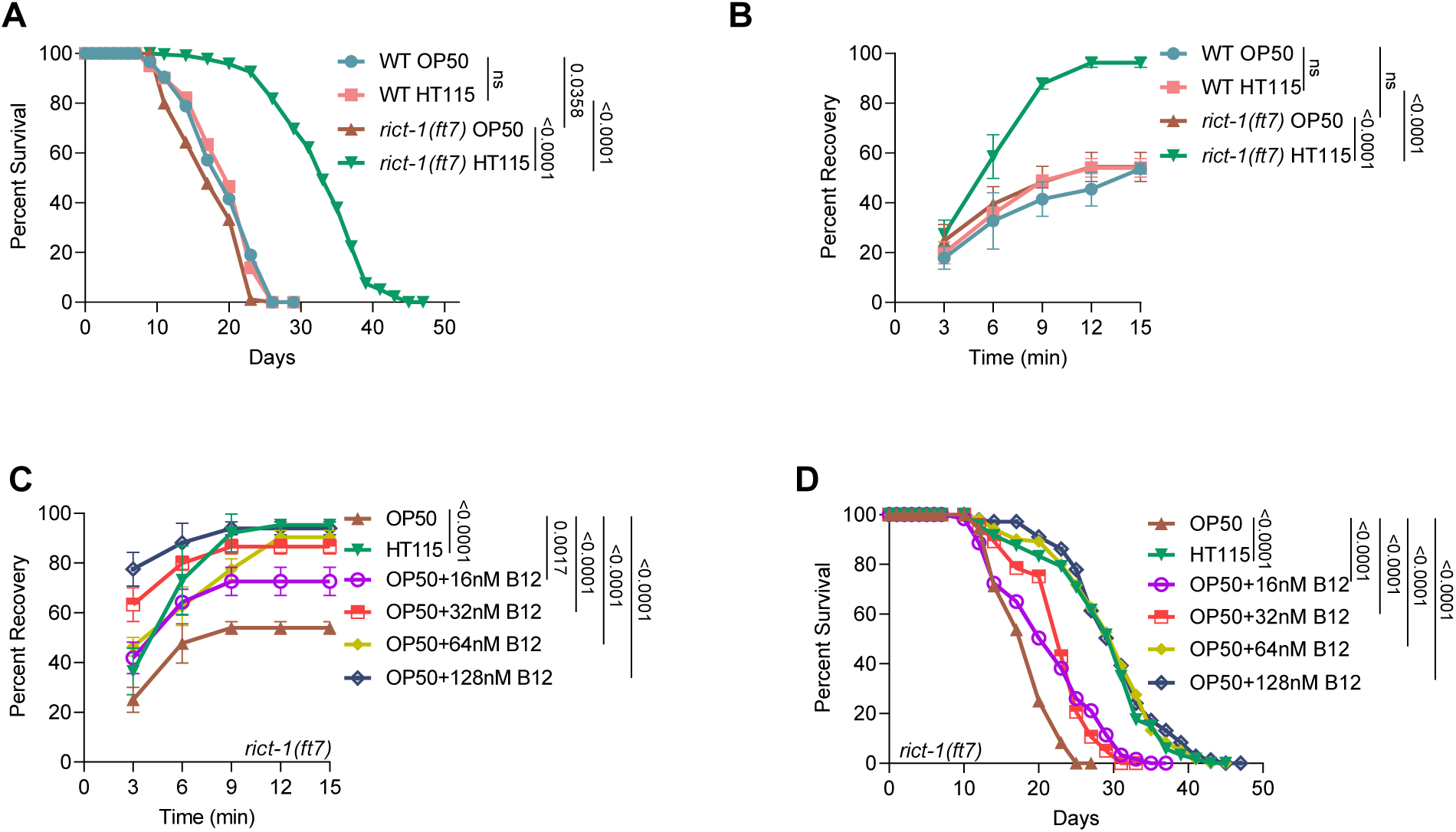
B12 is required for the osmotic stress tolerance (OST) and longevity of *rict-1(ft7)* worms. **(A)** Feeding *rict-1(ft7)* with HT115 resulted in a significant extension of lifespan compared to the mutant worms fed an OP50 diet. No effect on lifespan was observed in wild-type worms fed either diet. One of three biologically independent replicates is shown. P-value determined using Mann-Whitney U test. **(B)** The *rict-1(ft7)* worms demonstrated improved tolerance to osmotic stress when fed on HT115, relative to OP50. The wild-type worms were unaffected. One of three biologically independent replicates is shown. P-value determined using Ordinary Two-way ANOVA with Tukey’s multiple comparisons test. **(C)** The *rict-1(ft7)* worms displayed a dose-dependent increase in OST when grown on OP50 supplemented with B12. One of two biologically independent replicates is shown. P-value determined using Ordinary Two-way ANOVA with Tukey’s multiple comparisons test. **(D)** The *rict-1(ft7)* worms showed a dose-dependent increase in lifespan when fed a B12-supplemented OP50 diet. One of two biologically independent replicates is shown. P-value determined using Mann-Whitney U test. P-value ≥0.05 was considered not significant, ns. All experiments were performed at 20°C. All data and analysis are provided in the Source Data file.

Like mammals, *C. elegans* cannot synthesize B12 and entirely sources this micronutrient from the bacteria [21]. We and others have earlier demonstrated that the HT115 bacteria have higher B12 levels compared to OP50, and worms fed on a high B12 diet accumulate elevated levels of the micronutrient in the body [8]. So, we explored whether the diet-dependent changes in lifespan and osmotic stress tolerance in *rict-1(-)* can be explained by the difference in the micronutrient levels in the two bacterial strains. We supplemented OP50 with B12 and found that it restored the *rict-1* mutant’s osmotic stress tolerance (OST), in a concentration-dependent manner, to levels similar to those observed on the HT115 diet, while exerting minimal influence on the wild-type (**Figures 1C, S1A**). Additionally, B12 supplementation to the OP50 diet extended the lifespan of *rict-1(-)* worms, in a concentration-dependent manner, whereas wild-type worms remained unaffected (**Figures 1D, S1B**). Together, these data suggest that RICT-1 prevents alterations in OST and lifespan when the worms feed on a diverse bacterial diet containing varying levels of B12.

### The lifespan and osmotic stress tolerance of the *rict-1* mutant are dependent on Met-C of the host and the bacteria

In *C. elegans*, B12 is a cofactor for METR-1 (methionine synthase), which converts homocysteine (Hcy) to Met in Met-C. MTRR-1 (methionine synthase reductase) catalyses the NADPH-dependent reductive methylation of methionine synthase [22], maintaining the methionine synthase in an active state (**Figure 2A**). Since the *rict-1(-)* worms are dependent on B12 levels for increased OST and lifespan, we asked whether the mutant would also require functional *metr-1/mtrr-1.* We knocked down these genes by feeding the worms double-stranded RNA generated in HT115 (HT115i) or OP50 (OP50i) supplemented with B12. Interestingly, knocking down *metr-1* or *mtrr-1* in HT115i or OP50i supplemented with B12 abrogated the increased OST of *rict-1(-)* worms with little effect on wild-type (**Figures 2B, 2C, S2A**). On knocking down these two genes, the wild-type worms exhibited no change in lifespan, but the lifespan of *rict-1(-)* was suppressed (**Figures 2D, S2B**). This highlights the essential role of the functional Met-C in providing the OST and lifespan benefits to *rict-1(-)* worms on a high B12 diet.

**Figure 2:**
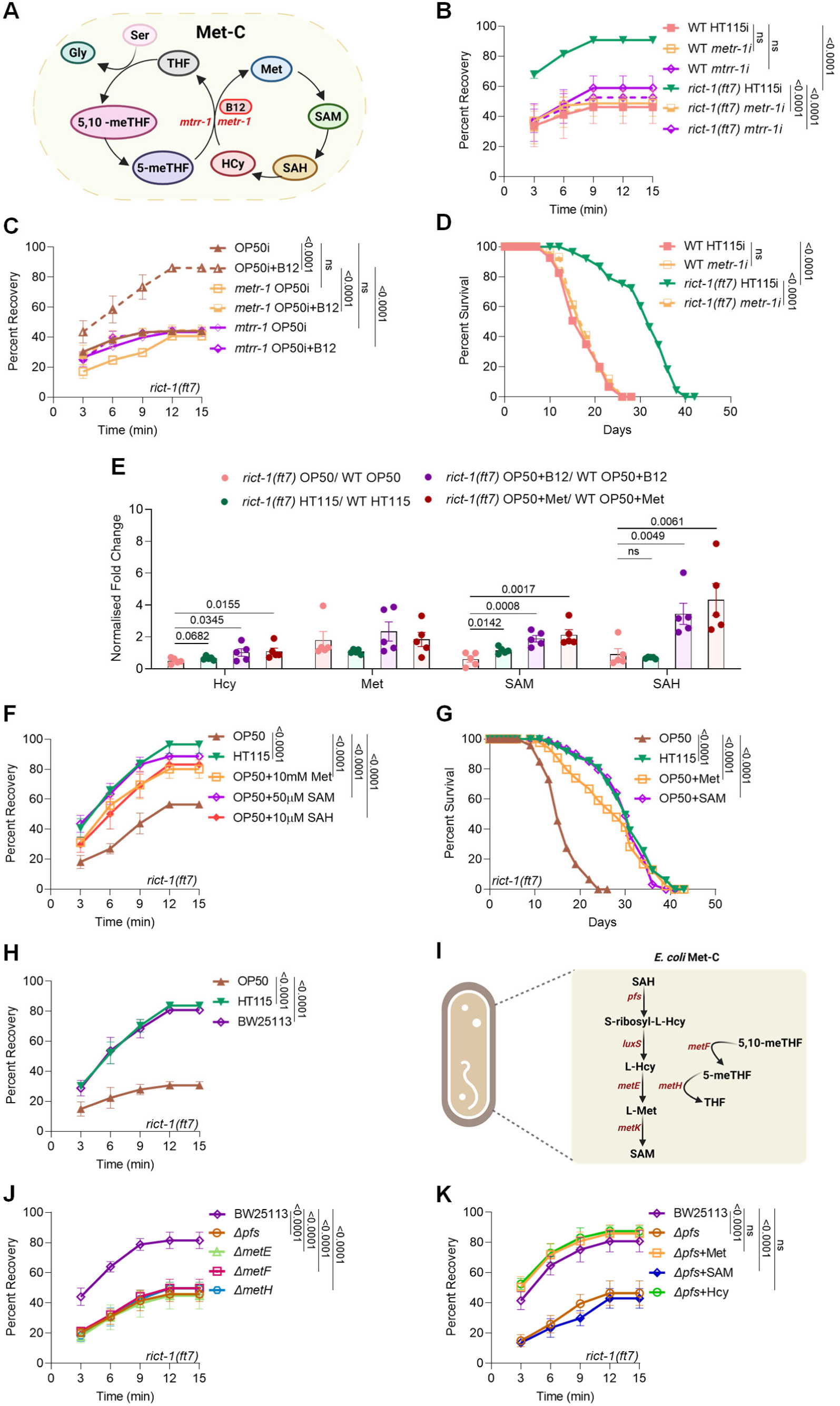
Host and bacterial Met-C modulate lifespan and osmotic stress tolerance of *rict-1(ft7).* **(A)** A simplified view of the one-carbon cycle (Met-C) in *C. elegans*. Methionine synthase (METR-1) requires B12 as a cofactor. Created in BioRender. **(B)** The increased OST of *rict-1(ft7)* was suppressed when *metr-1* or *mtrr-1* was knocked down using HT115i (RNAi using *E. coli* HT115). Wild-type worms remained unaffected by both RNAi. One of two biologically independent replicates is shown. P-value determined using Ordinary Two-way ANOVA with Tukey’s multiple comparisons test. **(C**) The increased OST observed for *rict-1(ft7)* grown on 64nM B12-supplemented control OP50i (RNAi using *E. coli* OP50) was suppressed on *metr*-*1* or *mtrr*-*1* was knocked down. One of three biologically independent replicates is shown. P-value determined using Ordinary Two-way ANOVA with Tukey’s multiple comparisons test. **(D)** A reduction in the lifespan of *rict-1(ft7)* worms was observed upon knockdown of *metr-1* using HT115i. One of two biologically independent replicates is shown. P-value determined using Mann-Whitney U test. **(E)** The relative abundance of Met-C intermediates, as determined by targeted metabolomics, in *rict-1(ft7)* versus wild-type worms grown on varying B12 or Met *E. coli* diet. Each data point represents a biologically independent replicate (n=5). Statistical differences were evaluated using a one-tailed unpaired t-test. **(F)** The OST of *rict-1(ft7)* worms was restored when the OP50 diet was supplemented with Met, SAM, or SAH, reaching levels similar to those observed with the HT115 diet. One of two biologically independent replicates is shown. P-value determined using Ordinary Two-way ANOVA with Tukey’s multiple comparisons test. **(G)** The lifespan of *rict-1(ft7)* worms was restored upon supplementation of Met or SAM to the OP50 diet, comparable to the mutants grown on HT115. One of two biologically independent replicates is shown. P-value determined using Mann-Whitney U test. **(H)** The *rict-1(ft7)* worms showed an increase in the OST when grown on BW25113, a B12-rich strain (parental strain of the Keio collection library). One of three biologically independent replicates is shown. P-value determined using Ordinary Two-way ANOVA with Tukey’s multiple comparisons test. **(I)** A schematic diagram showing key enzymes involved in bacterial Met-C. Created in BioRender. **(J)** The *rict-1(ft7)* worms showed a reduction in OST when grown on the deletion mutants of the *E. coli* Met-C genes, *pfs*, *metE, metF,* or *metH*. One of three biologically independent replicates is shown. P-value determined using Ordinary Two-way ANOVA with Tukey’s multiple comparisons test. **(K)** Supplementation of Met or Homocysteine (Hcy), but not SAM to *E. coli Δpfs* resulted in an increase in the OST of *rict-1(ft7)*. One of two biologically independent replicates is shown. P-value determined using Ordinary Two-way ANOVA with Tukey’s multiple comparisons test. P-value ≥0.05 was considered not significant, ns. All experiments were performed at 20 °C. All data and analysis are provided in the Source Data file.

Given the dependency of *rict-1(-)* worms on both B12 and the Met-C, and the above findings showing enhanced stress resilience and lifespan on a B12-rich HT115 diet, we hypothesized that the mutant may exhibit increased flux through the Met-C under high-B12 conditions. To test this, we performed targeted metabolomic profiling of key Met-C intermediates in wild-type and *rict-1(-)* worms grown on either the OP50 or HT115. We quantified the levels of Hcy, Met, S-adenosylmethionine (SAM), and S-adenosylhomocysteine (SAH), which serve as indicators of Met-C activity. In *rict-1(-)* worms fed with HT115 diet, we observed an increase in Hcy and SAM levels relative to wild-type controls on the same diet (**Figure 2E**). However, the relative ratio of Met and SAH in *rict-1(-)* and wild-type grown on either diet remained the same, possibly hinting at an increased utilisation of these metabolites. Next, we explored how supplementation with specific metabolic inputs, either B12 or Met, would influence the abundance of Met-C intermediates. Worms were grown on an OP50 alone or supplemented with B12 or Met, and metabolite levels were assessed in both wild-type and *rict-1(-)* worms. Supplementation with either B12 or Met led to a significant elevation of Hcy, SAM, and SAH, specifically in *rict-1(-)* worms (**Figure 2E**). In summary, the metabolic landscape of *rict-1* mutants is dynamically shaped by diet. By supplying either B12 or Met, we are able to unmask a latent capacity for enhanced Met-C throughput that is uniquely amplified in the absence of RICT-1 function.

Next, we asked whether the Met-C metabolites would rescue the reduced OST observed in the *rict-1(-)* fed on the OP50 diet. While the wild-type worms remained unaffected by the supplementation of Met, SAH, and Hcy, supplementation of SAM in OP50 causes a small increase in OST (**Figures S2C, S2D**). The addition of Met, SAM, SAH, or Hcy to the OP50 diet restored the OST of the *rict-1(-)* worms to levels comparable to those on the HT115 diet (**Figures 2F, S2D**). Further, when the wild-type worms were fed an OP50 diet supplemented with Met or SAM, it resulted in a small but significant increase in lifespan (**Figure S2E**). The *rict-1(-)* worms showed a dramatic increase in the lifespan when supplied with Met or SAM in OP50, similar to the levels on the HT115 diet (**Figure 2G**). These findings show that the *rict-1* mutant benefits from the higher flux through the Met-C of the host when fed a B12-rich diet.

Next, we investigated whether the bacterial Met-C exerts any influence on host physiology. To this end, we analyzed the OST in *C. elegans* when fed with different *E. coli* strains: OP50, HT115, and BW25113-the latter being a parental strain enriched in B12 [23, 24]. We found that *rict-1*(-) worms displayed a marked elevation in OST when grown on BW25113, reaching levels similar to those observed with the HT115 diet (**Figure 2H**). In contrast, wild-type worms showed no significant OST variation across these bacterial diets (**Figure S2F**).

Next, we grew both the wild-type and *rict-1(-)* worms on the strains of *E. coli* where key genes of its Met-C (*pfs, metE, metF* and *metH*) are deleted (**Figure 2I**), comparing the phenotypes with the parental strain BW25113. We found that the deletion of Met-C genes of the bacteria led to a reduction in the OST of *rict-1(-)* worms (**Figure 2J**), without any effect on wild-type (**Figure S2G)**.

To determine which bacterial metabolite is required for the altered OST of *C. elegans* grown on the high-B12 diet, we supplemented Met, SAM, and Hcy to the Δ*pfs* strain, where the first gene in the biosynthesis of Met is deleted (**Figure 2I**). Notably, the increase in the OST of *rict-1(-)* was observed only on the supplementation of Met or Hcy (which produces Met) but not SAM (**Figure 2K**); wild-type showed no difference in OST (**Figure S2H)**.

Together, these results highlight the significance of the essential amino acid Met as well as B12 originating from the bacteria, which in turn modulates the Met-C of *rict-1(-)* worms, ultimately impacting the host’s OST. The elevated B12 may assist both the bacterial as well as the host’s Met-C in generating and utilizing Met, respectively.

### Transsulfuration and propionate catabolic pathway enzymes are essential for osmotic stress tolerance and lifespan extension of the *rict-1* mutant

The Hcy generated in the Met-C is channelled into the transsulfuration pathway and is converted into succinyl-CoA through a multi-step process. The toxic odd-chain fatty acid oxidation product, propionate, is also detoxified through this pathway [25]. Importantly, a second B12-requiring mitochondrial enzyme, methylmalonyl-CoA mutase, encoded by *mmcm-1* in *C. elegans*, functions in this pathway [25]. A non-canonical B12-independent shunt pathway, gated by the rate-limiting enzyme acyl-CoA dehydrogenase (ACDH-1), can also detoxify propionate to produce acetyl-CoA [25] (**Figure 3A**). To evaluate the contribution of these pathways, we knocked down the key enzymes like *cbs-1* (cystathionine beta synthase), *pcca-1* (propionyl-CoA carboxylase), and *mce-1* (methylmalonyl-CoA epimerase), as well as *mmcm-1*, and assessed their impact on OST. Knocking down any of these genes reduced OST in *rict-1(-)* (**Figures 3B, 3C**), while knockdown of *cbs-1* or *mce-1* led to a small but significant increase in wild-type worms (**Figures 3C, S3A**). However, the shunt pathway is not involved in *rict-1(-)* OST on high B12 diet, as *acdh-1* knockdown had no effects (**Figure S3B**). Knocking down *mmcm-1* also reduced the enhanced lifespan of the *rict-1(-)* worms without significantly affecting wild-type (**Figure 3D**). These findings indicate that the longevity benefits of the *rict-1(-)* on high B12 diet are dependent on the transsulfuration pathway and the canonical propionate catabolic pathway, but not on the shunt pathway.

**Figure 3:**
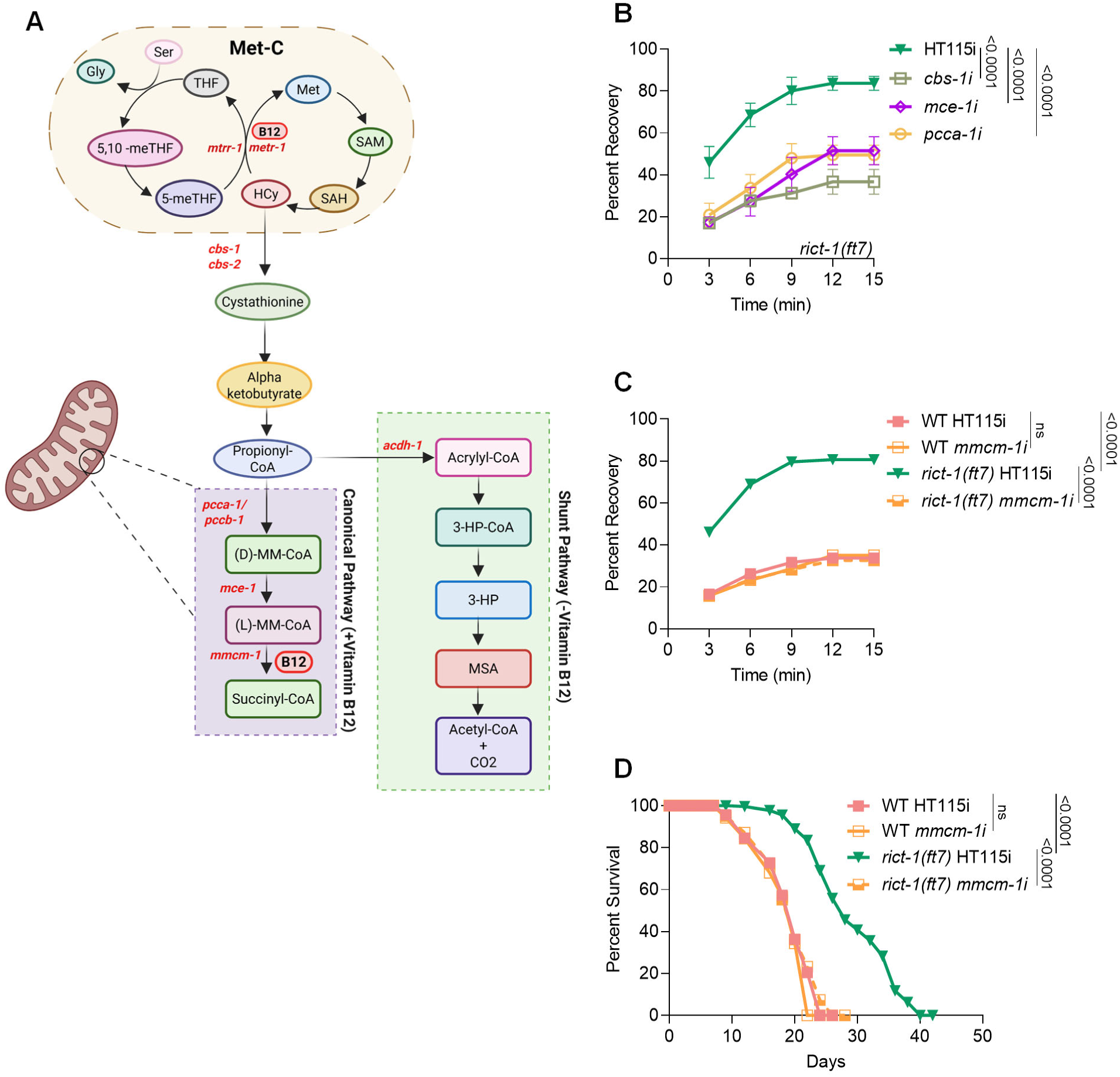
Osmotic stress tolerance of *rict-1(ft7)* is dependent on the transsulfuration and propionate catabolic pathway. **(A)** A schematic view of the B12-dependent metabolic pathways in *C. elegans.* Apart from the METR-1, the MMCM-1 (methylmalonyl-CoA mutase-1) uses B12 as a cofactor. In the presence of low levels of B12, the shunt pathway is used for propionate catabolism, where ACDH-1 (acyl-CoA dehydrogenase-1) is the rate-limiting enzyme. Created in BioRender. **(B)** Knockdown of *cbs-1* (cystathionine beta-synthase), *pcca-1* (propionyl coenzyme A carboxylase alpha subunit), and *mce-1* (methylmalonyl-CoA epimerase) using HT115i reduced the OST of s*rict-1(ft7)* worms. One of three biologically independent replicates is shown. P-value determined using Ordinary Two-way ANOVA with Tukey’s multiple comparisons test. **(C)** Knockdown of *mmcm-1* using HT115i resulted in a decrease in OST of the *rict-1(ft7)* worms, while wild-type worms remained unaffected. One of three biologically independent replicates is shown. P-value determined using Ordinary Two-way ANOVA with Tukey’s multiple comparisons test. **(D)** The *rict-1(ft7)* worms showed a reduction in lifespan on knockdown of *mmcm-1* using HT115i, while wild-type worms remained unaffected. One of three biologically independent replicates is shown. P-value determined using Mann-Whitney U test. P-value ≥0.05 was considered not significant, ns. All experiments were performed at 20 °C. All data and analysis are provided in the Source Data file.

### Increased levels of succinate provide osmotic stress tolerance and lifespan benefits to the *rict-1* mutant

Succinyl-CoA, generated through the B12-dependent canonical propionate catabolic pathway, enters the Krebs cycle and is converted to succinate by the action of succinyl-CoA synthetase (**Figure 4A**). In *C. elegans*, this enzyme has two subunits encoded by *suca-1* and *sucg-1*. We asked whether succinate produced from succinyl-CoA may be responsible for the increased stress tolerance of *rict-1(-)* worms. To investigate this, we knocked down *suca-1* or *sucg-1* using HT115i and assessed OST. We found that the knockdown of *suca-1* or *sucg-1* reduced the OST of *rict-1(-)* with no effect on wild-type (**Figures 4B, S4A**). Next, we asked whether other Krebs cycle genes are involved in the diet-dependent enhanced stress tolerance of *rict-1(-)* worms. For this, we silenced all genes associated with this pathway and conducted an end-point OST assay. Notably, the OST of *rict-1(-)* was unaffected by the gene knockdowns, except for *sucg-1/suca-1*, suggesting that the other Krebs cycle genes in general are not required by the *rict-1(-)* worms for enhanced OST (**Figure S4B**).

**Figure 4:**
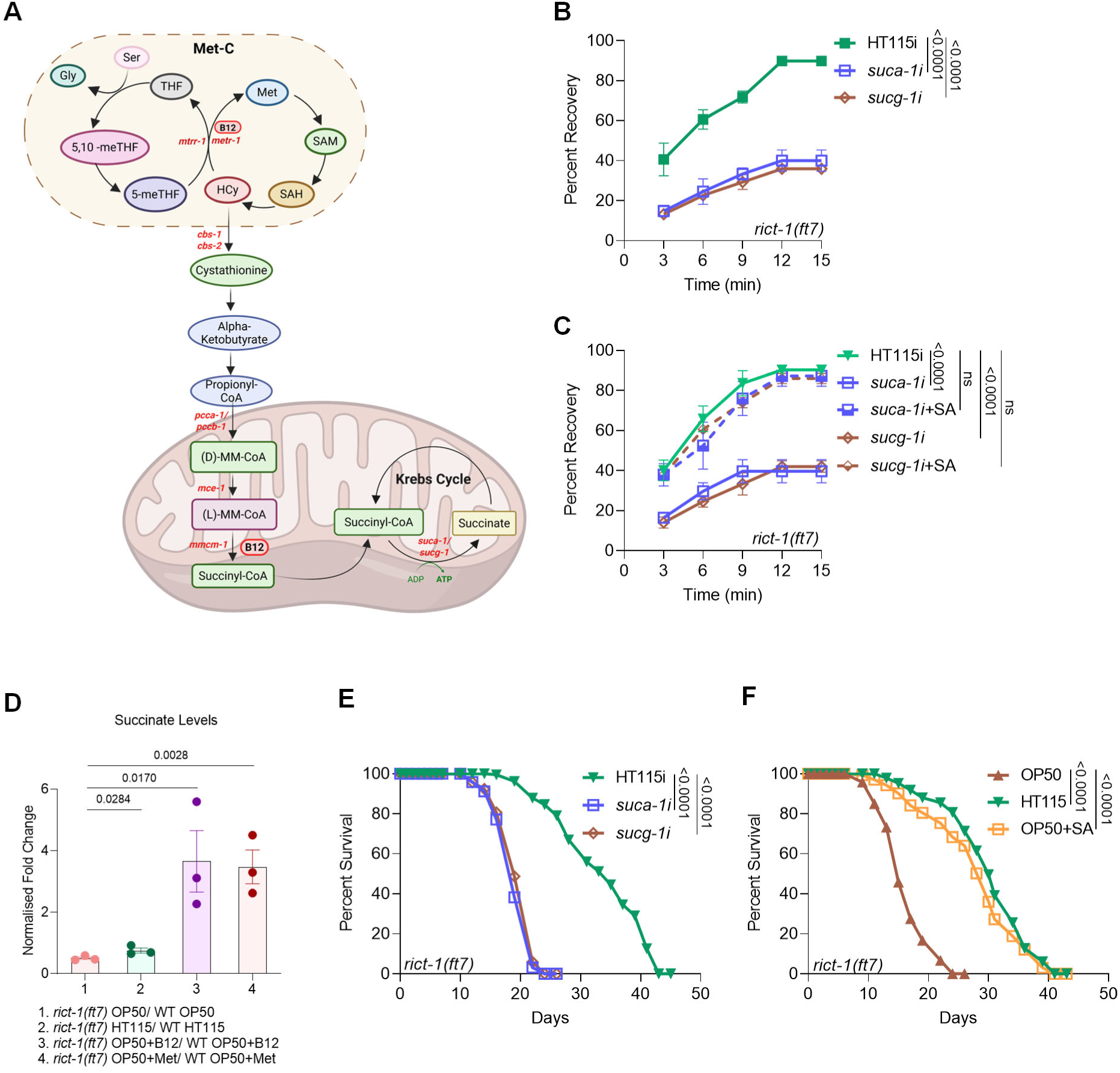
Elevated succinate levels confer osmotic stress resistance and promote longevity in *rict-1(ft7)*. **(A)** A schematic illustrating the generation of succinate from succinyl-CoA by the enzymatic activity of mitochondrial succinyl-CoA synthetase containing two subunits, *suca-1* and *sucg-1* in *C. elegans*. Created in BioRender. **(B)** The *rict-1(ft7)* worms showed a suppression in OST on knockdown of *suca-1* or *sucg-1* using HT115i. One of three biologically independent replicates is shown. P-value determined using Ordinary Two-way ANOVA with Tukey’s multiple comparisons test. **(C)** The OST of *rict-1(ft7)* grown on *suca-1* or *sucg-1* HT115i was restored on supplementation of SA. One of two biologically independent replicates is shown. P-value determined using Ordinary Two-way ANOVA with Tukey’s multiple comparisons test. **(D)** The relative abundance of succinate, as determined by targeted metabolomics, in *rict-1(ft7)* versus wild-type worms grown on varying B12 or Met *E. coli* diet. Each data point represents a biologically independent replicate (n=3). Statistical significance was assessed using a one-tailed t-test. **(E)** Lifespan of *rict-1(ft7)* worms showed a suppression upon knockdown of *suca-1* or *sucg-1* using HT115i. One of three biologically independent replicates is shown. P-value determined using Mann-Whitney U test. **(F)** Supplementation of SA to OP50 resulted in an increase in the lifespan of *rict-1(ft7)* worms. One of two biologically independent replicates is shown. P-value determined using Mann-Whitney U test. P-value ≥0.05 was considered not significant, ns. All experiments were performed at 20 °C. All data and analysis are provided in the Source Data file.

Further, we supplemented succinic acid (SA) after knocking down *suca-1* or *sucg-1* to determine whether succinate levels are crucial for the OST benefits observed in *rict-1(-)* worms. We found that supplementing SA rescued the OST in the *rict-1(-)* worms with *suca-1* or *sucg-1* knockdown, while wild-type showed no changes (**Figures 4C, S4C**), highlighting the importance of increased succinate levels for the stress tolerance phenotype. Supporting this notion, targeted metabolomics analysis via LC-MS revealed that HT115 diet or supplementation of OP50 with B12 or Met led to a substantial elevation in succinate levels in *rict-1(-)* worms compared to wild-type (**Figure 4D**). This difference was particularly pronounced when comparing mutant worms fed on B12 or Met-supplemented diet, suggesting that nutrient availability can amplify succinate accumulation in a *rict-1*-sensitive manner.

We next examined the effects of *suca-1* and *sucg-1* knockdown on worm lifespan. Knockdown of either gene led to a significant reduction in the lifespan of *rict-1(-)* worms (**Figure 4E**), whereas wild-type worms were largely unaffected (**Figure S4D**). Additionally, the supplementation of OP50 with SA extended the lifespan of *rict-1(-)* worms (**Figure 4F**) but had no impact on the lifespan of wild-type worms (**Figure S4E**).

These findings collectively support a model in which elevated succinate levels, either through endogenous biosynthesis or dietary supplementation, are sufficient to promote lifespan extension in the *rict-1(-)* worms. The absence of similar effects in wild-type worms implies a rewired metabolic state in *rict-1*-deficient worms that renders succinate a key longevity-supporting metabolite, potentially through its role in mitochondrial function or signalling pathways linked to metabolic stress adaptation.

### Altered mitochondrial dynamics in *rict-1* mutant fed HT115 diet

Given that *mmcm-1*, *sucg-1,* and *suca-1* function within the mitochondrial matrix and their loss disrupts stress resistance in *rict-1(-)*, we hypothesized that mitochondrial structure itself might be altered in a diet- and genotype-specific manner. To test this, we investigated mitochondrial morphology of L4 stage worms using the *zcIs14[myo-3p*::GFP(mit)] strain, which labels body wall muscle mitochondria with GFP, allowing *in vivo* visualization of mitochondrial networks via confocal microscopy.

Interestingly, striking morphological differences were observed across dietary conditions. The *rict-1(-)* worms exhibited a highly fragmented mitochondrial network when fed on the HT115 diet, indicative of increased mitochondrial fission or impaired fusion. In contrast, *rict-1(-)* worms maintained on the OP50 diet displayed a predominantly fused and elongated mitochondrial network, reflecting enhanced connectivity (**Figure 5A**). Notably, wild-type worms maintained a fused mitochondrial morphology under both dietary conditions, suggesting that the observed fragmentation is specific to the *rict-1* genotype in the context of the HT115 diet (**Figure 5A**).

**Figure 5:**
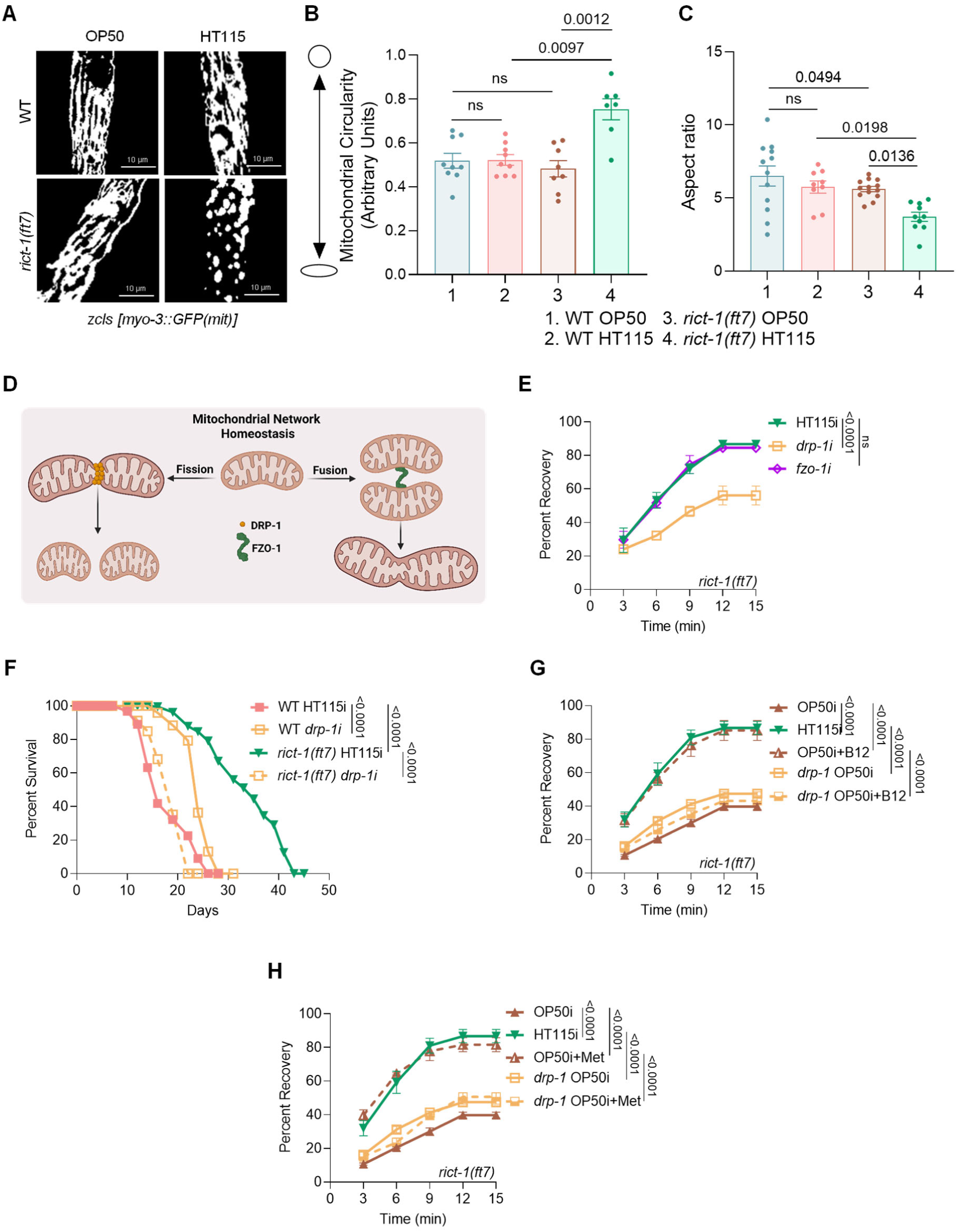
Diet-dependent alterations in mitochondrial dynamics in *rict-1(ft7)*. **(A)** The *rict-1(ft7)* worms showed mitochondrial fragmentation on the HT115 diet, while a fused mitochondrial network was observed on the OP50 diet. One of three biologically independent replicates is shown. **(B)** An increase in mitochondrial circularity was observed in *rict-1(ft7)* worms fed on HT115 as compared to the OP50 diet. **(C)** The Aspect ratio was reduced when *rict-1(ft7)* worms were grown on the HT115 diet. P-value determined using Ordinary Two-way ANOVA with Tukey’s multiple comparisons test. **(D)** A schematic representation of the mitochondrial dynamics showing the involvement of DRP-1 and FZO-1 in inducing mitochondrial fragmentation and fusion, respectively. Created in BioRender. **(E)** The *rict-1(ft7)* worms exhibited reduced OST on *drp-1* HT115i, which would reduce fragmentation of mitochondria. No change in OST is observed in *rict-1(ft7)* fed with *fzo-1* HT115i, which would increase mitochondrial fragmentation. One of three biologically independent replicates is shown. P-value determined using Ordinary Two-way ANOVA with Tukey’s multiple comparisons test. **(F)** Lifespan of *rict-1(ft7)* was reduced upon knockdown of *drp-1* as compared to control. One of three biologically independent replicates is shown. P-value determined using Mann-Whitney U test. **(G)** Supplementation of B12 failed to rescue the OST of *rict-1(ft7)* worms when *drp-1* was knocked down using OP50i. One of three biologically independent replicates is shown. P-value determined using Ordinary Two-way ANOVA with Tukey’s multiple comparisons test. **(H)** Supplementation of Met failed to rescue the OST of *rict-1(ft7)* worms when *drp-1* was knocked down using OP50i. One of three biologically independent replicates is shown. P-value determined using Ordinary Two-way ANOVA with Tukey’s multiple comparisons test. P-value ≥0.05 was considered not significant, ns. All experiments were performed at 20 °C. All data and analysis are provided in the Source Data file.

To quantitatively assess these structural differences, we employed morphometric analyses to calculate mitochondrial circularity and aspect ratio (major axis to minor axis length, which indicates the shape of mitochondria). Consistent with the visual observations, *rict-1(-)* worms grown on HT115 exhibited significantly higher mitochondrial circularity and a reduced aspect ratio, hallmarks of a fragmented network (**Figures 5B, 5C**). In contrast, *rict-1(-)* worms on OP50 and wild-type worms on either diet maintained lower circularity and higher aspect ratios, consistent with an elongated mitochondrial state (**Figures 5B, 5C**). This suggests a potential role of mitochondrial fission in stress tolerance and longevity of the *rict-1(-)* worms.

### Osmotic stress tolerance and lifespan of the *rict-1* mutant require fragmentated mitochondria

Subsequently, we investigated whether the fragmentation of mitochondria of *rict-1(-)* in the HT115 diet is the underlying cause or a consequence of heightened OST. To accomplish this, we silenced the expression of two proteins, FZO-1 and DRP-1, which regulate mitochondrial fusion and induce mitochondrial fragmentation, respectively (**Figure 5D**), and performed OST assay. Wild-type worms subjected to either *drp-1* or *fzo-1* RNAi resulted in a significant enhancement in OST (**Figure S5A**). This suggests that the knockdown of these genes positively impacts the ability of wild-type worms to withstand osmotic stress, possibly through mitohormesis [26]. However, a decrease in OST was specifically observed in the *rict-1(-)* when exposed to *drp-1* HT115i (**Figure 5E**), which is associated with reduced fragmentation of mitochondria. On the other hand, no change was observed in *rict-1(-)* worms fed with *fzo-1* HT115i (**Figure 5E**), which would typically result in heightened mitochondrial fragmentation. The *drp-1* RNAi dramatically reduced the lifespan of *rict-1(-)* while it increased the lifespan of wild-type (**Figure 5F**). In line with this result, B12 or Met supplementation to the OP50i diet failed to increase OST if *drp-1* was knocked down in the *rict-1(-)* worms (**Figures 5G, 5H**). Wild-type worms, as usual, showed no changes in either *drp-1* or *fzo-1* OP50i on supplementation of B12 or Met (**Figures S5B, S5C, S5D, S5E**). However, heightened OST of *rict-1(-)* does not require FZO-1 under the same circumstances (**Figures S5F, S5G**). This underscores the crucial involvement of the DRP-1 protein, and consequently, mitochondrial fission, in supporting the augmented OST and lifespan of *rict-1(-)*.

### Met-C regulates mitochondrial fragmentation in the *rict-1* mutant fed a high B12 or Met diet

We have found that wild-type worms have a highly connected mitochondrial network irrespective of diet, while the *rict-1(-)* worms exhibit fragmented mitochondria on the HT115 diet. We have also established the requirement of proteins involved in the regulation of mitochondrial dynamics in OST and the lifespan of the *rict-1(-)* worms. So next, we asked whether the metabolites and genes of the Met-C also regulate the mitochondrial dynamics in the mutant. We found that when B12 was added to the OP50 diet, mitochondria in *rict-1(-)* worms appeared highly fragmented and punctate, with a clear disruption of the elongated network. Quantification revealed increased mitochondrial circularity and a concomitant reduction in aspect ratio, indicating enhanced fission activity (**Figures 6A, 6B, 6C**). Similarly, Met supplementation also led to mitochondrial fragmentation in *rict-1(-)* worms, phenocopying the effects of B12 (**Figures 6D, 6E, 6F**). In contrast, in the wild-type worms, supplementation with either B12 or Met had minimal effect on mitochondrial morphology, with predominantly tubular and interconnected networks, similar to that on OP50 (**Figures 6A, 6B, 6C, 6D, 6E, 6F**). This indicates that enhanced Met-C flux does not appreciably influence mitochondrial morphology in the wild-type worms.

**Figure 6:**
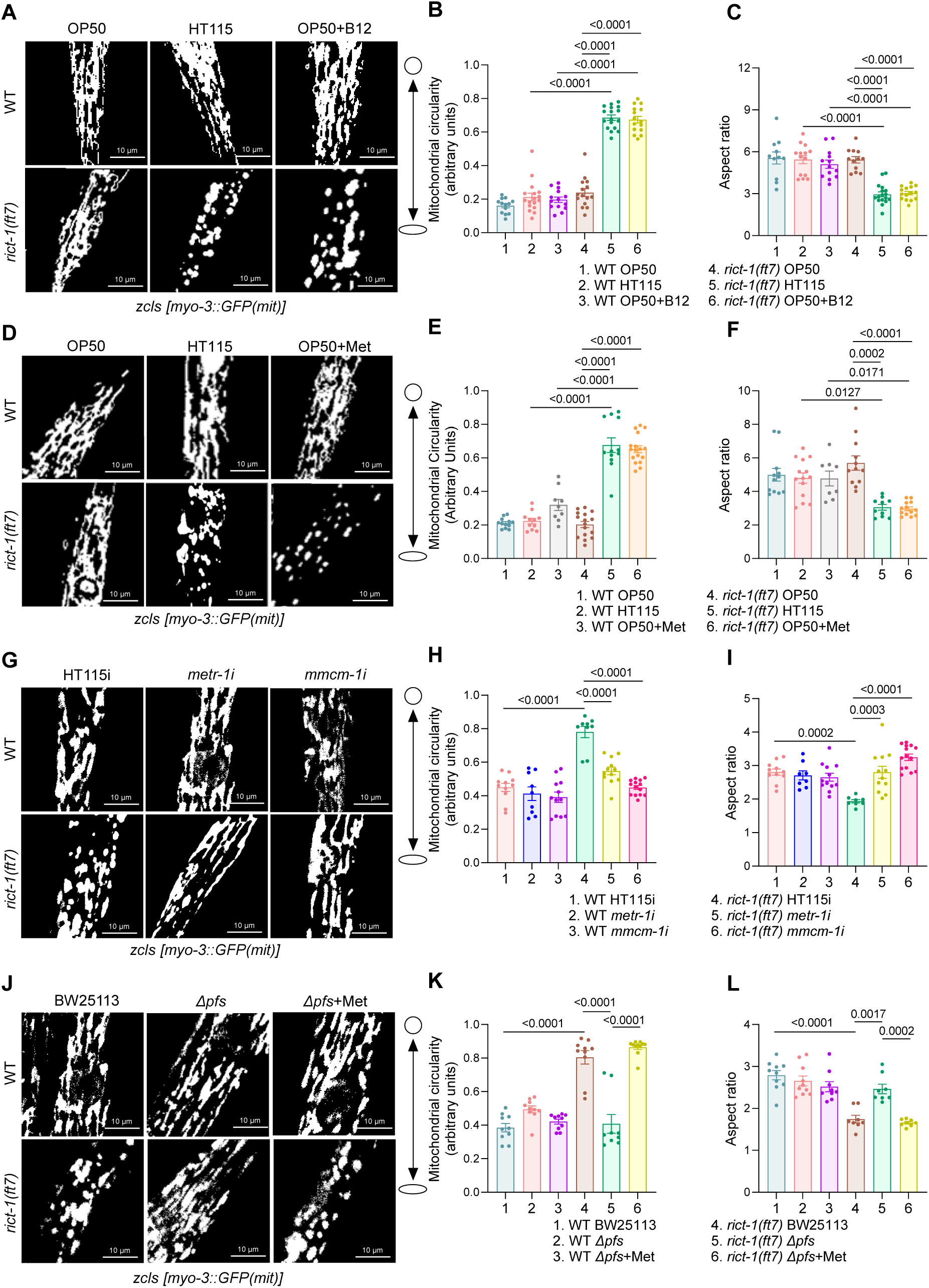
Host and bacterial Met-C modulates mitochondrial dynamics in *rict-1(ft7)*. **(A-C)** Supplementation of B12 to OP50 induced fragmentation of mitochondria in *rict-1(ft7)*, similar to that on HT115 diet. (A) Representative images. (B) Mitochondrial circularity increases and (C) aspect ratio reduces in *rict-1(ft7)* worms grown on high B12 diet. **(D-F)** Supplementation of Met to OP50 induced fragmentation of mitochondria in *rict-1(ft7)*, similar to that on HT115 diet. (D) Representative images. (E) Mitochondrial circularity increases and (F) aspect ratio reduces in *rict-1(ft7)* worms grown on Met-supplemented OP50 diet. **(G-I)** The *rict-1(ft7)* worms showed a fused mitochondrial morphology upon knockdown of *metr-1* or *mmcm-1* using HT115i. (G) Representative images. (H) Mitochondrial circularity reduces and (I) aspect ratio increases on knockdown of *metr-1* or *mmcm-1* using HT115i in *rict-1(ft7)*. **(J-L)** Feeding *E. coli Δpfs* showed a fused mitochondrial morphology in *rict-1(ft7)* worms, which was reversed on the addition of Met. (J) Representative images. (K) Mitochondrial circularity increases and (L) aspect ratio reduces in *rict-1(ft7)* worms when *Δpfs* is supplemented with Met. One of two biologically independent replicates is shown for all the experiments. P-value determined using Ordinary Two-way ANOVA with Tukey’s multiple comparisons test. P-value ≥0.05 was considered not significant, ns. All experiments were performed at 20 °C. All data and analysis are provided in the Source Data file.

Further, we focused on the Met-C and the mitochondrial propionate catabolism arm, both of which are critically dependent on B12 availability and represent central hubs for coordinating methylation potential and metabolic homeostasis. RNAi-mediated depletion of *metr-1* or *mmcm-1* resulted in a striking reversal of mitochondrial fragmentation in the *rict-1(-)* worms, revealing a transition from punctate and fragmented mitochondria to a more elongated, tubular network (**Figures 6G, 6H, 6I**). This highlights the crucial role of the host’s Met-C metabolism in regulating mitochondrial morphology in the *rict-1(-)* worms, suggesting that the OST and lifespan benefits are closely linked to mitochondrial architecture.

Next, to assess whether the bacterial Met-C contributes to host mitochondrial architecture, we examined the impact of microbial Met-C functionality on mitochondrial morphology in the *rict-1(-)* worms. We utilized the *E. coli* Δ*pfs* strain, comparing it with the parental BW25113. We found that when *rict-1(-)* worms were fed the Δ*pfs* strain, mitochondrial networks exhibited a pronounced fused and elongated phenotype, in contrast to the fragmented morphology typically observed when the mutant is maintained on B12-rich BW25113 (**Figures 6J, 6K, 6L**). Importantly, the mitochondrial fragmentation in the mutant could be rescued by dietary Met supplementation, which restored the fragmented mitochondrial phenotype (**Figures 6J, 6K, 6L**). These findings indicate that bacterial Met-C output is both necessary and sufficient to influence mitochondrial dynamics in the host when mTORC2 signalling is compromised.

### Succinate drives mitochondrial fragmentation in the *rict-1* mutant

Given that OST in *rict-1(-)* depends on the succinyl-CoA synthetase subunits, SUCA-1 and SUCG-1, we postulated that elevated succinate levels may promote mitochondrial fragmentation, thereby enhancing OST and lifespan. To test this hypothesis, we performed RNAi-mediated knockdown of *suca-1* and *sucg-1* in *rict-1(-)* worms and assessed mitochondrial morphology. Knockdown of *suca-1* or *sucg-1* resulted in a significant reduction of mitochondrial fragmentation, revealing a shift towards more elongated and fused mitochondrial networks in *rict-1(-)* worms (**Figure 7A, 7B, 7C, 7D, 7E, 7F**).

**Figure 7:**
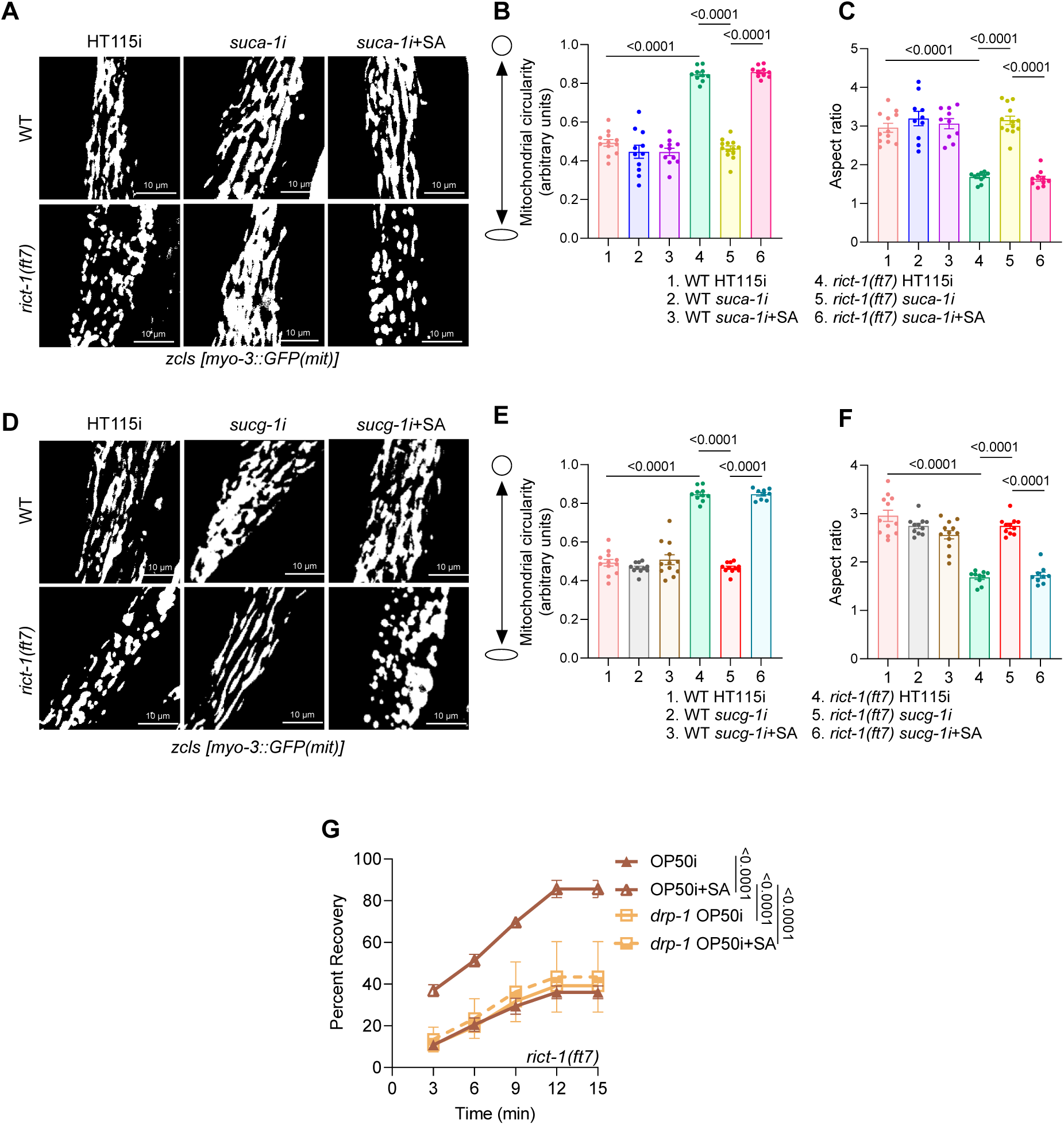
Succinate promotes mitochondrial fragmentation in *rict-1(ft7)*. **(A-C)** The *rict-1(ft7)* worms showed fused mitochondrial morphology upon knockdown of *suca-1* using HT115i, which was reversed on supplementation of SA. (A) Representative images. (B) Mitochondrial circularity increases and (C) aspect ratio reduces in *rict-1(ft7)* on the addition of SA in the *suca-1i*. **(D-F)** The *rict-1(ft7)* worms showed fused mitochondrial morphology upon knockdown of *sucg-1* using HT115i, which was reversed on supplementation of SA. (A) Representative images. (B) Mitochondrial circularity increases and (C) aspect ratio reduces in *rict-1(ft7)* on addition of SA in the *sucg-1i*. One of two biologically independent replicates is shown for all the experiments. **(G)** Supplementation of SA to OP50i showed an increase in OST in the *rict-1(ft7)* worms. However, no significant changes were observed in *rict-1(ft7)* worms when SA was supplemented along with the *drp-1* OP50i. One of three biologically independent replicates is shown. P-value determined using Ordinary Two-way ANOVA with Tukey’s multiple comparisons test. P-value ≥0.05 was considered not significant, ns. All experiments were performed at 20 °C. All data and analysis are provided in the Source Data file.

To further validate the role of succinate, we supplemented SA concurrently with *suca-1* or *sucg-1* RNAi and examined mitochondrial morphology. We found that when SA supplementation restored mitochondrial fragmentation, demonstrated by increased circularity and decreased aspect ratio compared to knockdown alone in *rict-1(-)* worms (**Figures 7A, 7B, 7C, 7D, 7E, 7F**). In contrast, wild-type worms consistently exhibited a fused mitochondrial network across all tested conditions (**Figures 7A, 7B, 7C, 7D, 7E, 7F**). Furthermore, as expected, the *rict-1(-)* fed on OP50 supplemented with SA exhibited a fragmented mitochondrial phenotype comparable to that observed on the HT115 diet, while wild-type worms maintained fused mitochondrial networks under all tested conditions (**Figures S6A, S6B, S6C**). These findings indicate that succinate acts as a metabolic signal driving mitochondrial fragmentation specifically in the *rict-1* mutant, linking succinate accumulation to altered mitochondrial dynamics and enhanced stress tolerance.

To determine whether mitochondrial fragmentation is a necessary downstream effector of succinate-induced physiological benefits in *rict-1(-)* worms, we interrogated the requirement for DRP-1-dependent mitochondrial fission in this context. We found that wild-type worms displayed no significant alterations in OST upon SA supplementation, either in the presence or absence of *drp-1*(RNAi) (**Figure S6D**). Interestingly, despite the presence of exogenous succinate, *drp-1* HT115i led to a substantial attenuation of OST in *rict-1(-)* worms (**Figure 7G**). These results suggest that succinate alone is not sufficient to confer enhanced OST to the *rict-1(-)* worms in the absence of DRP-1-mediated mitochondrial fission, pointing to a hierarchical relationship where succinate acts upstream to induce a fission-dependent cellular state that promotes stress tolerance.

### Mitophagy induced in *rict-1* mutant is dependent on a Met-C-succinate axis

Having established that DRP-1-dependent mitochondrial fragmentation is required for succinate-mediated stress tolerance in *rict-1(-)* worms, we next investigated whether this remodelling promotes mitophagy. Mitochondrial fission often precedes mitophagy and is thought to facilitate the segregation and removal of dysfunctional mitochondrial fragments, thus preserving mitochondrial homeostasis [27]. Increased mitophagy has been shown to delay aging and increase lifespan [28], and in the context of the *rict-1* mutant, could potentially connect its metabolic rewiring to the enhanced stress tolerance and longevity.

To assess mitophagy, we employed the *C. elegans* MitoRosella biosensor strain, which expresses a fragment of mitochondrial outer membrane protein TOMM-20 fused to a pH-stable DsRed and a pH-sensitive GFP [29]. In this system, progression of mitochondria into acidic autolysosomal compartments quenches GFP fluorescence while preserving the DsRed signal, enabling quantification of mitophagy via the GFP:DsRed fluorescence ratio.

The *rict-1(-)* worms showed robust mitochondrial fragmentation when fed with HT115 diet, resulting in a significant decrease in the GFP:DsRed ratio compared to worms fed with the OP50 diet (**Figure 8A**). This indicates enhanced mitophagy, specifically on the HT115 diet. In contrast, wild-type worms exhibited no significant diet-dependent changes in mitophagy, consistent with their fused mitochondrial morphology across dietary conditions (**Figure 8A**). This mitochondrial turnover likely acts as a key effector mechanism linking dietary quality to cellular homeostasis and organismal fitness in RICTOR-deficient worms.

**Figure 8:**
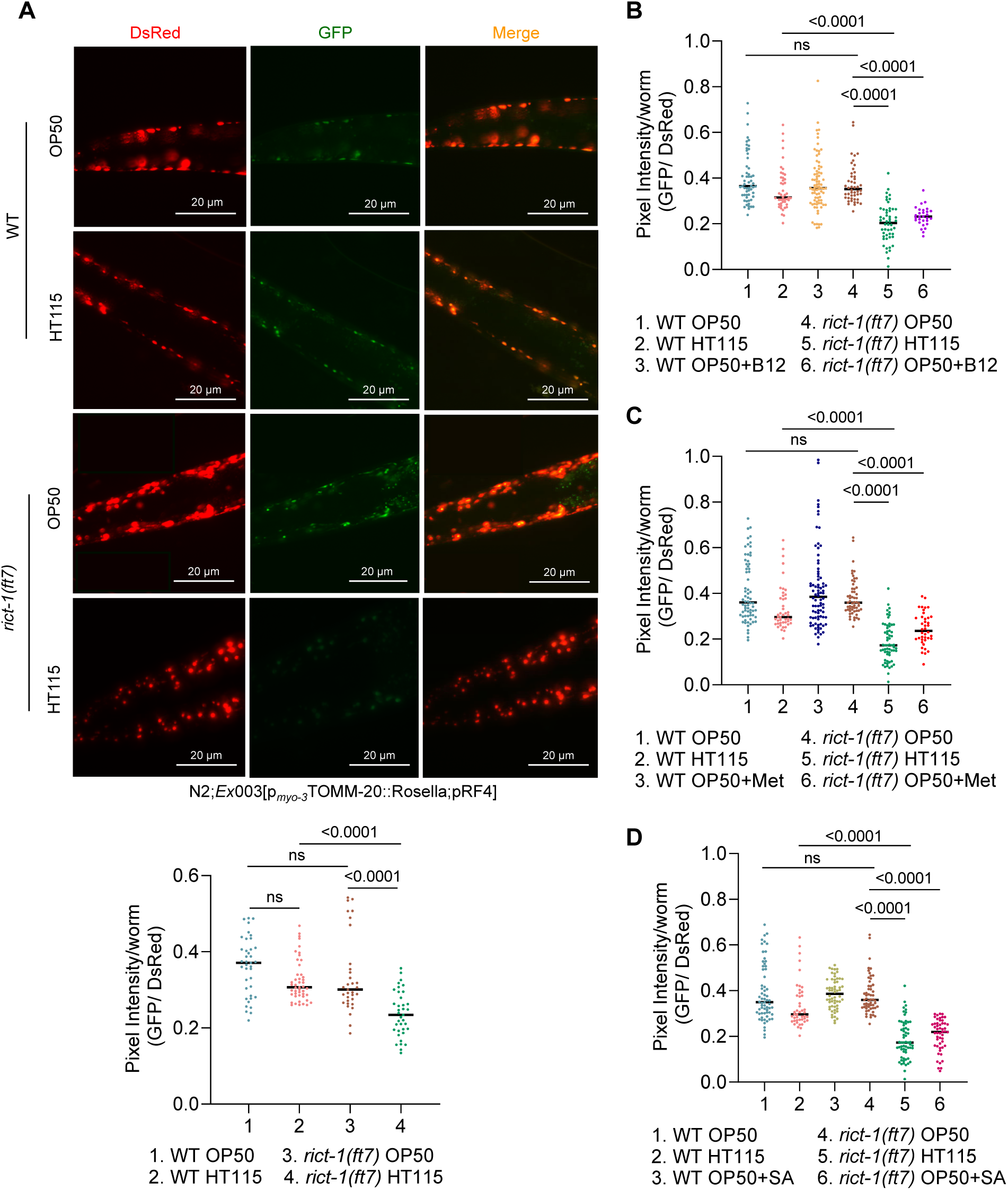
B12-, Met- and SA-rich diets induce mitophagy in *rict-1(ft7)*. **(A)** The HT115-fed *rict-1(ft7)* worms showed an increase in mitophagy, indicated by the reduced GFP/DsRed, when compared to the OP50 diet. Wild-type worms showed no change. Images captured at 40X magnification. (A) One of three biologically independent replicates is shown in the representative images. Quantification, provided below the images, shows compiled data of three biologically independent replicates. **(B-D)** Quantified data of *rict-1(ft7)* worms fed on (B) B12-supplemented, (C) Met-supplemented, or (D) SA-supplemented OP50 diet showed an increase in mitophagy, indicated by the reduced GFP:DsRed ratio, similar to the HT115 diet. Wild-type worms showed no change. Compiled data of three biologically independent replicates are shown. P-value determined using Ordinary Two-way ANOVA with Tukey’s multiple comparisons test. All experiments were performed at 20 °C. All data and analysis are provided in the Source Data file.

To interrogate the role of dietary metabolites in modulating mitochondrial quality control, we evaluated the effects of specific Met-C intermediates on mitophagy in *rict-1(-)* worms. Supplementation of the OP50 diet with B12 or Met resulted in a robust induction of mitophagy in the *rict-1(-)* worms, closely phenocopying the mitophagy-enhancing effects of the HT115 bacterial diet (**Figures 8B, 8C, S7A**).

In contrast, wild-type worms exhibited consistently lower mitophagy across all dietary conditions (**Figures 8B, 8C, S7B**), revealing a unique metabolic sensitivity in the *rict-1* mutant. These results support a model in which the altered metabolic state of *rict-1* mutants renders them responsive to Met-C perturbations, triggering mitophagy as a pro-longevity adaptation.

Next, we examined whether elevated succinate levels, previously implicated in mitochondrial remodelling, are sufficient to trigger mitophagy in the context of *rict-1* loss-of-function. We found that OP50 supplemented with SA led to a pronounced increase in mitophagy in the *rict-1(-)* worms, comparable to that observed on the HT115 diet (**Figures 8D, S7C**) while wild-type worms showed no changes on either diet (**Figure S7D**), highlighting the specificity of the succinate response in the *rict-1* mutant background.

These findings support the concept that succinate is not merely a downstream metabolic intermediate but acts as a signalling metabolite that activates mitophagy in a context-dependent manner. In *rict-1(-)* worms, succinate accumulation, either from endogenous metabolic rewiring or dietary supplementation, may serve as a bioenergetics and redox cue to initiate mitophagy, thereby contributing to enhanced mitochondrial turnover, stress tolerance, and longevity.

### Mitophagy is required for the increased osmotic stress tolerance and lifespan of *rict-1* mutant

Next, we investigated whether the increased mitochondrial fragmentation observed in *rict-1(-)* serves as an upstream trigger for mitophagy or if it is a consequence of mitophagy. For that, we knocked down *pink-1* or *pdr-1* (orthologues of mammalian PINK1 and PARKIN), two essential regulators of mitophagy initiation and progression [30] (**Figure 9A**), and assessed their effect on mitochondrial morphology. Even after knocking down *pink-1* or *pdr-1* in the *rict-1(-)* worms, they still retained a highly fragmented mitochondrial phenotype, as indicated by an elevated mitochondrial circularity and a reduced aspect ratio (**Figure 9B**). In contrast, wild-type worms subjected to the same RNAi treatments exhibited largely fused and tubular mitochondrial morphology, highlighting the specificity of this phenotype to the *rict-1* mutant context (**Figure 9B**).

**Figure 9:**
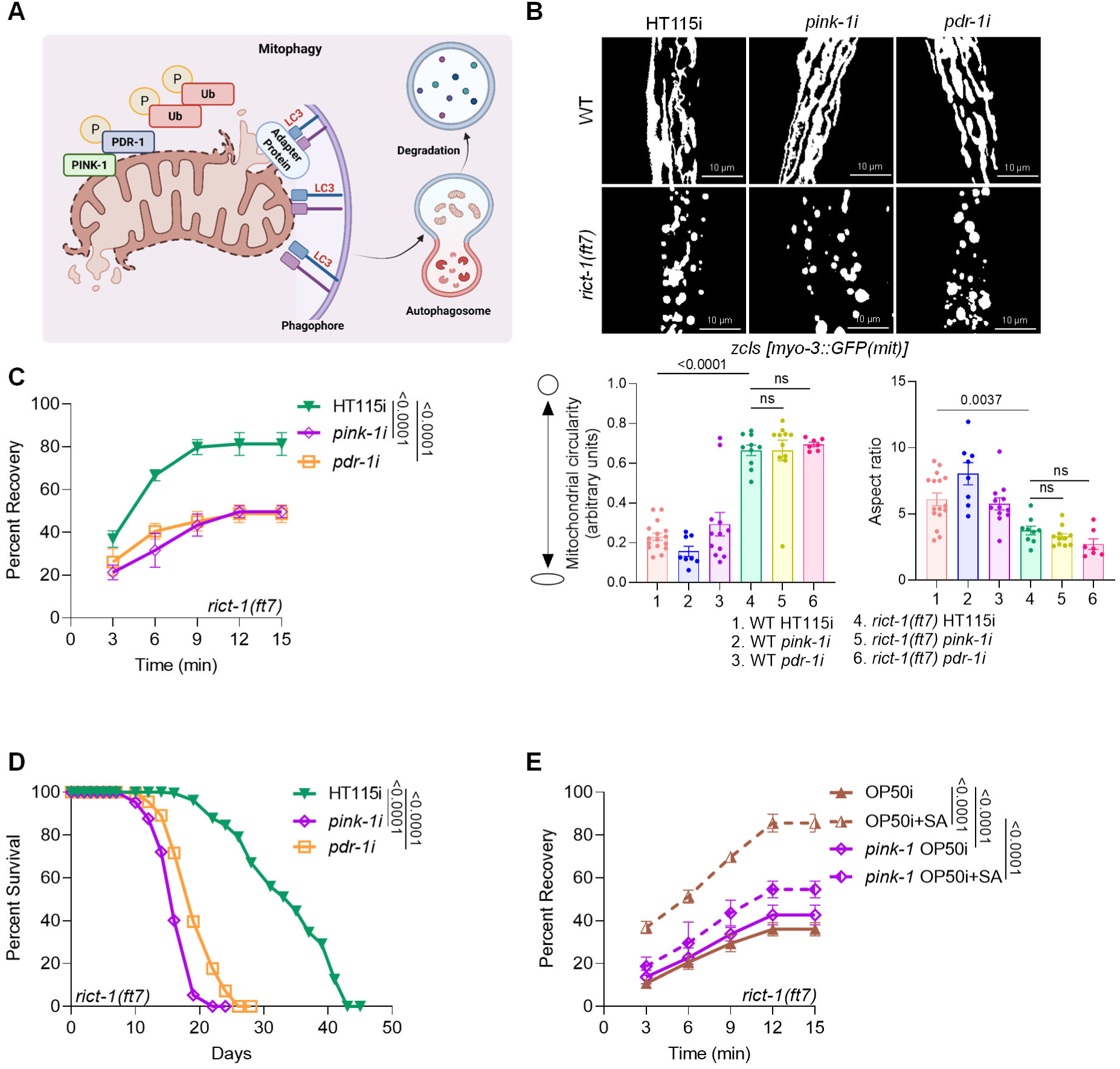
Mitophagy is essential for enhanced osmotic stress tolerance and lifespan of *rict-1(ft7)*. **(A)** A schematic representation of the mitophagy process highlighting the involvement of PINK-1 and PDR-1 proteins. Created in BioRender. **(B)** Representative images showing mitochondrial morphology in *rict-1(ft7)* worms following *pink-1* and *pdr-1* knockdown using HT115i. (Below) Compared to the control RNAi, mitochondrial circularity and the aspect ratio of the *rict-1(ft7)* worms remained unchanged when *pink-1* or *pdr-1* were knocked down. One of two biologically independent replicates is shown. P-value determined using Ordinary Two-way ANOVA with Tukey’s multiple comparisons test. **(C)** The *rict-1(ft7)* worms showed a reduction in OST on *pink-1* or *pdr-1* HT115i. One of three biologically independent replicates is shown. P-value determined using Ordinary Two-way ANOVA with Tukey’s multiple comparisons test. **(D)** The lifespan of *rict-1(ft7)* worms was suppressed upon knockdown of *pink-1* or *pdr-1* using HT115i. One of three biologically independent replicates is shown. P-value determined using Mann-Whitney U test. **(E)** Supplementation of SA following *pink-1* knockdown using OP50i showed no effect on OST of the *rict-1(ft7)* worms. One of three biologically independent replicates is shown. P-value determined using Ordinary Two-way ANOVA with Tukey’s multiple comparisons test. P-value ≥0.05 was considered not significant, ns. All experiments were performed at 20 °C. All data and analysis are provided in the Source Data file.

We next sought to determine whether mitophagy is causally linked to stress tolerance and longevity phenotypes characteristic of RICTOR-deficient worms. To test this, we knocked down *pink-1* or *pdr-1*, in both *rict-1(-)* and wild-type worms. RNAi-mediated depletion of either gene led to a significant reduction in OST in *rict-1(-)* worms, when fed the HT115 diet (**Figure 9C**). This phenotype was specific to the *rict-1* mutant background, as wild-type worms did not exhibit notable changes in OST under the same experimental conditions (**Figure S8A**).

Furthermore, suppression of mitophagy through *pink-1* or *pdr-1* knockdown significantly shortened the lifespan of *rict-1(-)* worms (**Figure 9D**), but also led to a slight reduction in the wild-type worms (**Figure S8B**), reinforcing the notion that the cellular phenomenon for mitochondrial turnover is essential for the maintenance of longevity [31].

Previous results showed that elevated succinate levels promote mitochondrial fragmentation in the *rict-1* mutant. This raised the question of whether succinate-induced physiological benefits depend on the activation of mitophagy machinery. To explore this, we knocked down *pink-1* using RNAi (in OP50 background) in the *rict-1(-)* worms fed on a diet supplemented with SA. Interestingly, this combined treatment failed to enhance OST to the levels seen in OP50 supplemented with SA, indicating that SA supplementation was ineffective when *pink-1* function was compromised (**Figure 9E**). The wild-type worms showed no response to SA under any condition (**Figure S8C**). These data suggest that the protective effect of succinate is dependent upon an intact mitophagy pathway.

## Discussion

*C. elegans* serves as a simplified and genetically tractable model to dissect how dietary inputs influence mitochondrial homeostasis, stress resilience, and lifespan, offering insights that may be relevant to human health [32–35]. Our study shows that the mTORC2 complex protein RICTOR regulates adaptive capacity to a diet containing higher levels of B12 or Met. In the *rict-1* deletion mutant, stress tolerance and lifespan are increased only on a high B12 or Met diet. We show that the mutant grown on high B12 or Met diet engages the host Met-C and the canonical propionate catabolic pathway, thereby increasing the levels of succinate. Increased succinate fragments the mitochondria, activating the pro-longevity process of mitophagy (**Figure 10**). Since the wild-type worms are refractory to these changes, we posit that RICTOR/mTORC2 has evolved to buffer against abrupt changes to dietary B12 or Met, which is known to have dramatic effects on life history traits [36, 37].

**Figure 10:**
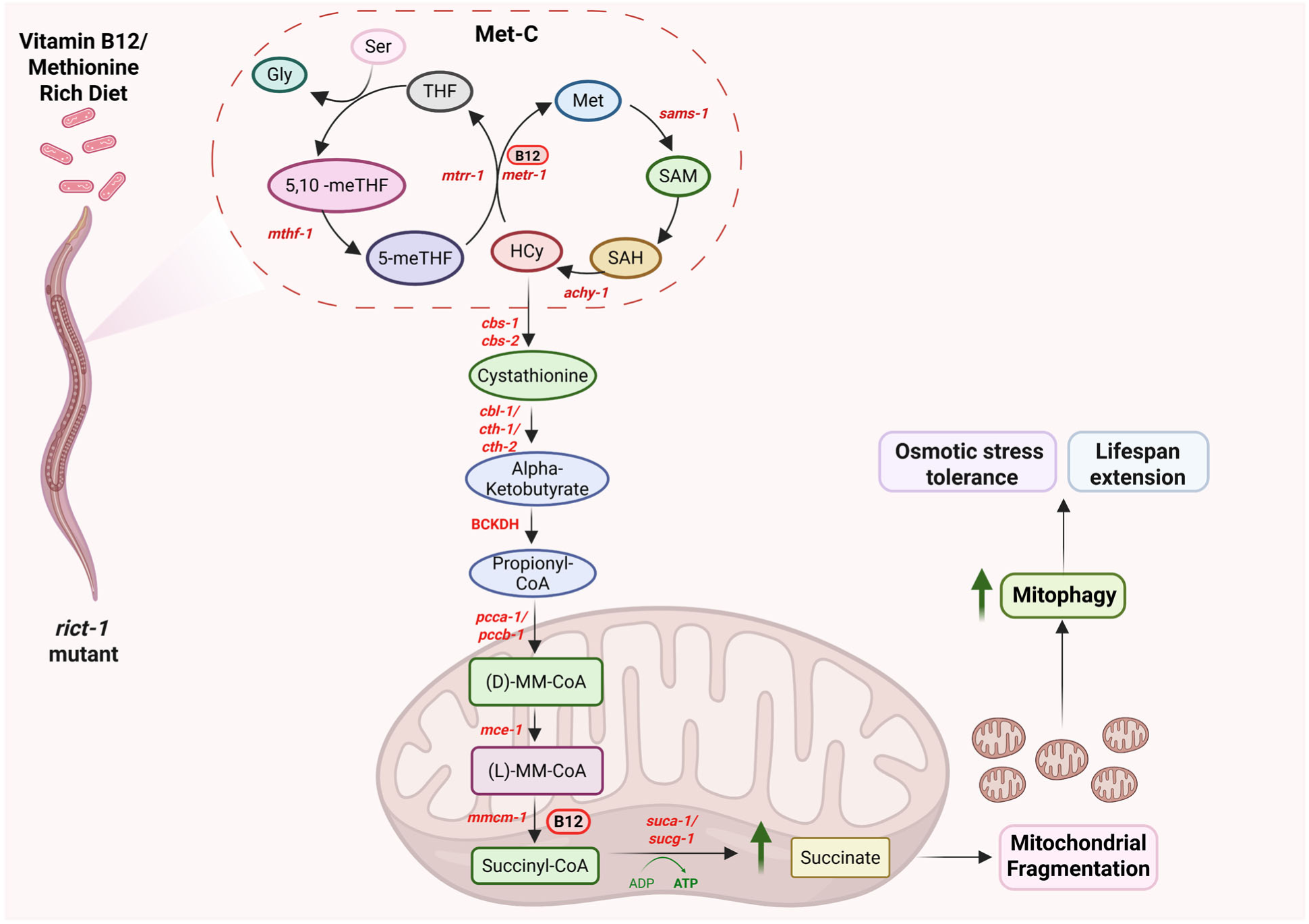
A model depicting RICT-1 regulation of longevity through a Met-C-mitophagy axis. The *rict-1(ft7)* worms fed on B12 or Met-rich diets display an increased osmotic stress tolerance and lifespan. Mechanistically, in the absence of RICT-1, the flux through the Met-C increases, resulting in an increased production of Met-C intermediates. This, in turn, leads to enhanced flow of Hcy into the transsulfuration pathway. This metabolic rerouting boosts levels of succinyl-CoA and succinate. Altogether, this results in fragmented mitochondria and enhanced mitophagy, ultimately contributing to osmotic stress tolerance and longevity benefits. Created in BioRender.

Although the mTOR pathway has been extensively studied for its regulation of growth and metabolic homeostasis in response to macronutrients [38], the role of mTORC2 in micronutrient sensing remains largely unexplored. Recent findings in mammalian systems suggest that mTORC1 and mTORC2 are sensitive to alterations to cellular redox states and methylation capacity [39–41], both of which are profoundly influenced by B vitamins such as B12 and folate [42, 43]. For example, folate deficiency has been shown to impair mTORC1 and mTORC2 activity by reducing phosphorylation of AKT in human trophoblasts, underscoring a conserved link between Met-C and mTOR signalling [44, 45].

Considering the central role of Met-C in cellular physiology and aging across species [46], this metabolic pathway is under tight regulation. For example, restriction of Met intake has been consistently shown to extend lifespan in various model organisms, including yeast, flies, and rodents, largely through the modulation of oxidative stress, mitochondrial activity, and the activation of AMPK signalling and autophagy [46–49]. Conversely, excess Met has been linked to elevated Hcy levels, vascular dysfunction, and increased risk of age-associated diseases such as cancer and cardiovascular disorders [50]. Together with our data, this suggests that the balance of Met-C is a key determinant of health and longevity, which is regulated by RICTOR/mTORC2.

B12 is a key cofactor in the conversion of methylmalonyl-CoA to succinyl-CoA, which is subsequently converted to succinate by the TCA cycle [51]. B12 deficiency leads to the accumulation of toxic intermediates and causes mitochondrial stress, as observed in disorders like methylmalonic acidemia [51]. Metabolites such as succinate have emerged as an important driver of mitochondrial fragmentation, mitophagy, and immune activation in both invertebrate and mammalian systems [52–54]. Succinate is known to alter mitochondrial dynamics [55, 56], linking metabolic imbalances to disease pathology [57]. Our identification of succinate-driven mitochondrial remodelling in *C. elegans* extends these concepts into the realm of diet-dependent lifespan regulation by micronutrients like B12.

The essential role of mitophagy in maintaining mitochondrial health is well-documented in the context of neurodegenerative diseases such as Parkinson’s and Alzheimer’s, where impaired clearance of damaged mitochondria contributes to neuronal dysfunction and degeneration [58]. On the other hand, enhancing mitophagy through genetic or pharmacological means has been shown to extend lifespan and improve stress tolerance in various model organisms [28, 59]. These findings collectively highlight the importance of safeguarding mitochondrial health during the aging process for promoting health and lifespan [60]. We show that in the *rict-1* mutant, bacterially derived B12 and Met orchestrate an adaptive mitochondrial remodelling response, where the mitochondria are fragmented and mitophagy is upregulated in early adult life. This may confer long-term benefits by preserving mitochondrial quality of the *rict-1* mutant in later stages of life.

Metabolic diseases such as diabetes and non-alcoholic fatty liver disease are being increasingly linked to mitochondrial dysfunction and altered nutrient sensing through pathways such as mTOR [61]. In this context, B12 deficiency has been implicated in exacerbating inflammation and oxidative stress, contributing to lifestyle disease progression [62, 63]. Our study shows how RICTOR/mTORC2 modulates host mitochondrial quality control machinery in response to bacterially derived nutrients, providing a valuable framework for developing dietary or pharmacological interventions aimed at improving metabolic resilience and delaying aging-related pathologies.

## Materials and Methods

### *C. elegans* Strains and Maintenance

*Caenorhabditis elegans* strains were acquired from the Caenorhabditis Genetics Center and maintained at 20°C on Nematode Growth Media (NGM) agar plates seeded with *Escherichia coli* OP50. All experiments were performed using L1 synchronized worms. The strains used include N2 Bristol as the wild-type, *rict-1(ft7)*, SJ4103 - *zcIs14* [*myo-3::GFP(mit)*], *rict-1(ft7)*;*zcIs14* [*myo-3::GFP(mit)*], IR1631: N2;*Ex003* [*p_myo-3_*TOMM-20::Rosella*; pRF4*] and *rict-1(ft7);*IR1631.

### Bacterial Growth

For bacterial cultures, glycerol stocks were streaked onto Luria-Bertani (LB) plates and incubated overnight (16 hours) at 37°C until single colonies were visible. A single colony was then transferred to LB broth and grown overnight (12 hours) at 37°C to establish the primary culture. Secondary cultures were grown by inoculating 1/100th of the primary culture and were incubated at 37°C until the optical density (OD at 600nm) reached 0.6. These secondary cultures were concentrated 10 times. A 350 µL and 1 mL aliquot of the concentrated cultures were plated on 60 mm and 90 mm NGM plates, respectively. After seeding, the plates were dried and incubated overnight to develop bacterial lawns. The bacterial strains used include *E. coli* OP50, HT115, and Keio collection clones [24] BW25113, *Δpfs, ΔmetE*, *ΔmetF*, and *ΔmetH*.

### RNAi Feeding

RNAi-feeding bacteria were sourced from the Ahringer or Vidal *C. elegans* RNAi library. Bacterial clones from glycerol stocks were streaked onto LB agar supplemented with Ampicillin (100 mg/L) and Tetracycline (12.5 mg/L), and incubated overnight (16 hours) at 37°C. Single colonies were then inoculated in LB broth containing ampicillin (100 µg/ml) and tetracycline (12.5 µg/ml) and grown overnight (12 hours) at 37°C to obtain the primary culture. For secondary cultures, a 1/100 dilution of the primary culture was incubated at 37°C until the OD600 reached 0.6. The cultures were then concentrated 10 times using M9 buffer with ampicillin (100 µg/ml) and IPTG (1 mM). A final volume of 350 µL and 1 mL cultures were plated on 60 mm and 90 mm NGM agar plates, respectively, supplemented with ampicillin (100 µg/ml) and IPTG (2 mM).

### Synchronization of Worms

Bleaching was performed to release eggs from the gravid worms grown on standard *E. coli* OP50. Further, they kept on rotation at 20°C for 16 hours for starvation. Then, the L1 worms were centrifuged at 3000 rpm for one minute, and the supernatant was decanted. The worms were poured onto the required experimental plates.

### Lifespan Assay

Synchronized population of L1 worms was cultured on NGM agar plates seeded with the required bacteria. At the late L4 stage, worms were transferred to plates overlaid with 5-fluorodeoxyuridine (FUDR; final concentration of 100 µg/ml). Lifespan assessments began on the 7th day of adulthood and were conducted every alternate day. Worms were scored as alive or dead by gently tapping with a platinum wire. Survival curves were analysed using the Mann-Whitney test (p-value at 50%) with the OASIS 2 software (https://sbi.postech.ac.kr/oasis2/).

### Osmotic Stress Assay

L4-staged worms grown on 50 mM NaCl containing NGM plates seeded with required bacterial feeds (supplemented or RNAi), were transferred to unseeded NGM plates containing 350 mM sodium chloride for 10 minutes. Followed by the recovery period, where the worms were transferred back to unseeded NGM plates (50 mM NaCl). The recovered worms were assessed over a time interval of 15 minutes to determine the percentage of recovery. GraphPad Prism 10 software was used to analyse the data. The 12th-minute survival values were used for reporting the statistics in the figures.

### Mitochondrial Morphology Analysis

Mitochondrial structures were assessed at the L4 larval stage using the SJ4103 strain, which harbours a GFP reporter specifically targeted to the mitochondrial matrix within body wall muscle cells. For microscopy, worms were immobilized with 20 mM tetramisole (N=3, n≥10) and then placed on 2% agarose-coated slides for imaging. Fluorescent images were acquired using a LSM980 microscope (Carl Zeiss, Germany), employing a 63X objective lens and using fluorescence excitation at 488 nm with emission collected at 520 nm. To ensure experimental consistency, imaging was performed on the first body wall muscle cell adjacent to the tail region. Mitochondrial morphology was quantitatively analysed in two ways: (1) mitochondrial circularity and (2) aspect ratio using the mitochondrial morphology analyser plugin within Fiji/ImageJ.

### Metabolites Supplementation

A 3 mM stock solution of Vitamin B12 (#V6629; Sigma Aldrich, USA) was prepared in the M9 buffer. A 1 mM working solution was generated by diluting the stock with 1X M9 buffer, followed by filtration using a 0.2-micron filter. Secondary *E. coli* cultures were concentrated 10-fold in 1X M9 buffer, and varying volumes of the 1 μM Vitamin B12 solution were added to achieve final concentrations of 16, 32, 64, or 128 nM in the bacterial feed. Thereafter, we used 64 nM for all experiments.

Methionine (#M9625; Sigma Aldrich, USA) was prepared as a 300 mM stock solution in Milli-Q water and filtered using a 0.2-micron filter. A 10-fold concentrated *E. coli* cultures in 1X M9 buffer were supplemented with appropriate volumes of the stock to yield final methionine concentrations of 1 (for mitochondrial morphology), 2.5, or 10 mM (OST, lifespan, and mitophagy).

S-adenosylmethionine (SAM; #A7007; Sigma Aldrich, USA) was dissolved in Milli-Q water to obtain a 1 M stock solution, which was filtered through a 0.2-micron filter. This stock was added to concentrated bacterial cultures to achieve a final concentration of 50 μM SAM in the feed.

S-adenosylhomocysteine (SAH; #A9384; Sigma Aldrich, USA) was prepared as a 1 mM stock solution in Milli-Q water and filtered through a 0.2-micron filter. A defined volume was added to the bacterial pellet to reach a final concentration of 10 μM SAH in the feed.

L-homocysteine (#69453; Sigma Aldrich, USA) was prepared as a 100 mM stock in Milli-Q water, filtered using a 0.2-micron filter, and added to the bacterial cultures to obtain a final concentration of 2.5 mM in the feed.

Succinic acid (#S3674; Sigma, USA) was dissolved in Milli-Q water to a concentration of 500 mM and filtered using a 0.2-micron filter. Appropriate volumes of the stock solution were added to the concentrated bacterial cultures to achieve final concentration of 2.5 mM.

### Mitophagy Imaging

Transgenic *C. elegans,* MitoRosella worms [IR1631 or *rict-1(ft7);*IR1631], were immobilized in a 20 mM tetramisole solution prepared in M9 buffer and transferred onto a 2% agarose pad. Individual animals were carefully placed in the droplet using a platinum wire loop. A coverslip was gently applied to secure the specimens for imaging. Fluorescent microscopy was carried out using an AxioImager M2 microscope (Carl Zeiss, Germany) at a magnification of 10X. Whole-worm fluorescent images were acquired for MitoRosella-expressing strains. All imaging parameters, including lens magnification, exposure times, filter sets, laser settings, and gain, were kept consistent throughout the experiments. Representative images were taken at 40X magnification.

Images were processed using ImageJ software. For intensity analysis, fluorescence from the whole worm was quantified. Channels were split, and regions of interest were manually outlined using the freehand selection tool. The area was measured using the “Measure” function. The ratio of GFP to DsRed fluorescence was calculated for each sample to assess relative signal levels. Statistical analyses were performed by using GraphPad Prism 10 software.

### Metabolomics

L4-staged worms of WT and *rict-1(ft7)* were grown under different conditions and collected in microcentrifuge tubes, washed with water thrice, and pelleted at 2000 rpm at 20 °C for 1 minute. Metabolites were extracted using a pre-chilled solvent consisting of Acetonitrile-methanol-water (3:5:2). For the extraction of metabolites, glass beads (#G8772, Sigma-Aldrich) equivalent to worm pellets were added to all the samples, respectively, followed by 30 seconds of vortexing and 30 seconds on ice. This vortexing step was repeated six times. Centrifugation was done at 15,000 × *g* at 4 °C for 15 minutes. The supernatant was collected in a fresh tube and vacuum dried. Followed by reconstitution in 25 μl of water with 0.1% formic acid. Centrifugation was performed at 15,000 x *g* for 10 minutes. 5 μl of sample was injected for LC-MS/MS analysis. The data acquisition was done using a UHPLC system coupled with a triple quadrupole mass spectrometer (TSQ Altis; Thermo) in a positive mode. 95% to 20% of buffer A was applied as a linear mobile phase. The gradient program was set as follows: 95% buffer B (1.5 min), 80%–50% buffer B (next 0.5 min), followed by 50% buffer B (next 2 min), and then decreased to 20% buffer B (next 50 s), 20% buffer B (next 2 min), and finally again 95% buffer B (next 4 min). Ion source parameters were used as follows: ion spray voltage at 3500 V, sheath gas at 45 Arbitrary unit (AU), auxiliary gas at 10 AU, ion transfer tube temperature at 290 °C and vaporizer temperature at 300 °C. Quantification was done using Multiple Reaction Monitoring (MRM) transitions. The method cycle was kept at 1 second. Five biological replicates were used, and the area was normalised to the protein amount. Skyline software was used to the relative quantification of metabolites. Fold change analysis was performed in MS Excel using one-tailed unpaired t-test.

### Quantification and statistical analysis

All statistical methods used in the paper are described in the figure legends, and additional details are provided in the methods section. Statistics were computed using GraphPad Prism 10. Survival curves were analyzed using the OASIS 2 software (https://sbi.postech.ac.kr/oasis2/).

## Acknowledgements

We thank the present and former members of the Molecular Aging Laboratory (National Institute of Immunology) for their support, Suhail Khan, and Smriti Raina in particular. The MitoRosella biosensor strain was a kind gift from Dr. Nektarios Tavernarakis. Some strains were provided by the *Caenorhabditis* Genetics Center (CGC), which is funded by the National Institutes of Health (NIH) Office of Research Infrastructure Programs (P40 OD010440). We thank the National Bioresource Project (NBRP), Japan, for the *E. coli* Keio collection deletion library. We thank the Central Confocal Microscopy facility and the Central Instrumentation facility of NII. Schematics in Figures 2A, 2H, 3A, 4A, 5D, 9A, and 10 were created with BioRender.

## Author contributions

AM and SM conceptualized the project. SM and SB conducted *C. elegans* experiments and analyzed data. SC and RU performed metabolomics under the supervision of SSG. AM, SM, and SB wrote the manuscript. AM supervised the project and acquired funding.

## Data availability statement

The authors declare that the main data supporting the findings of this study are available within the article and its supplementary files. The data reported in this manuscript are available in the Source data file that accompanies the manuscript.

