## Supplementary Figures for "RICTOR regulates an interspecies crosstalk that influences longevity through a novel methionine cycle-mitophagy axis"

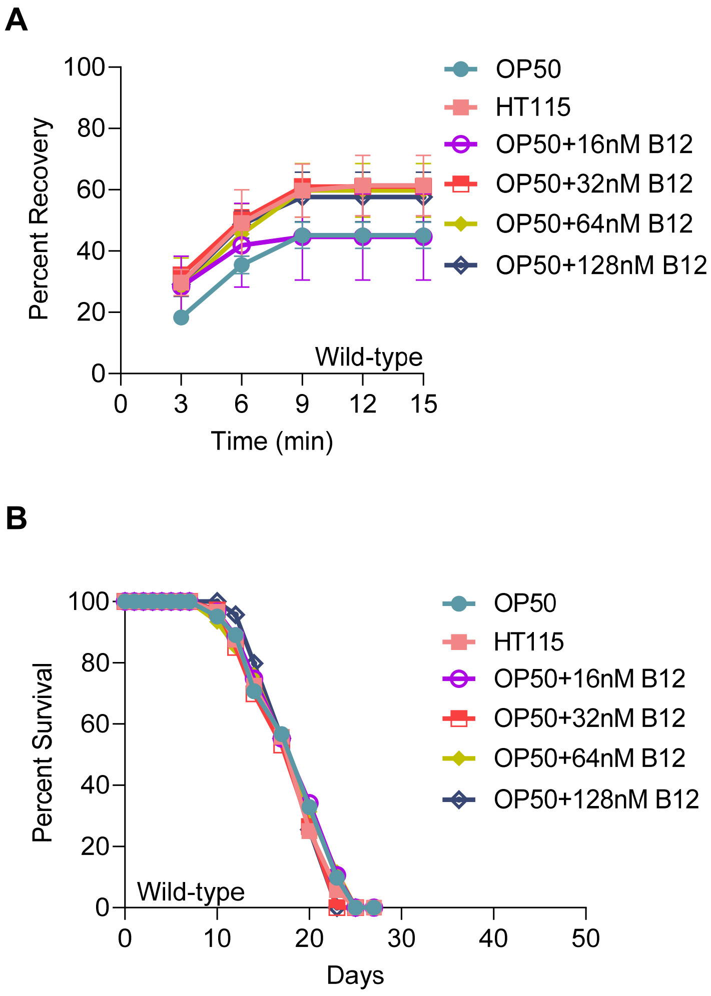


**Figure S1 (A)** No difference was observed in the OST of wild-type worms when grown on OP50 supplemented with B12. One of two biologically independent replicates is shown. P-value determined using Ordinary Two-way ANOVA with Tukey's multiple comparisons test and found to be non-significant. **(B)** The lifespan of wild-type worms remained unaffected by feeding B12-supplemented OP50. One of two biologically independent replicates is shown. P-value determined using Mann-Whitney U test and found to be non-significant. All experiments were performed at 20 $^{o}$C. All data and analysis are provided in the Source Data file.

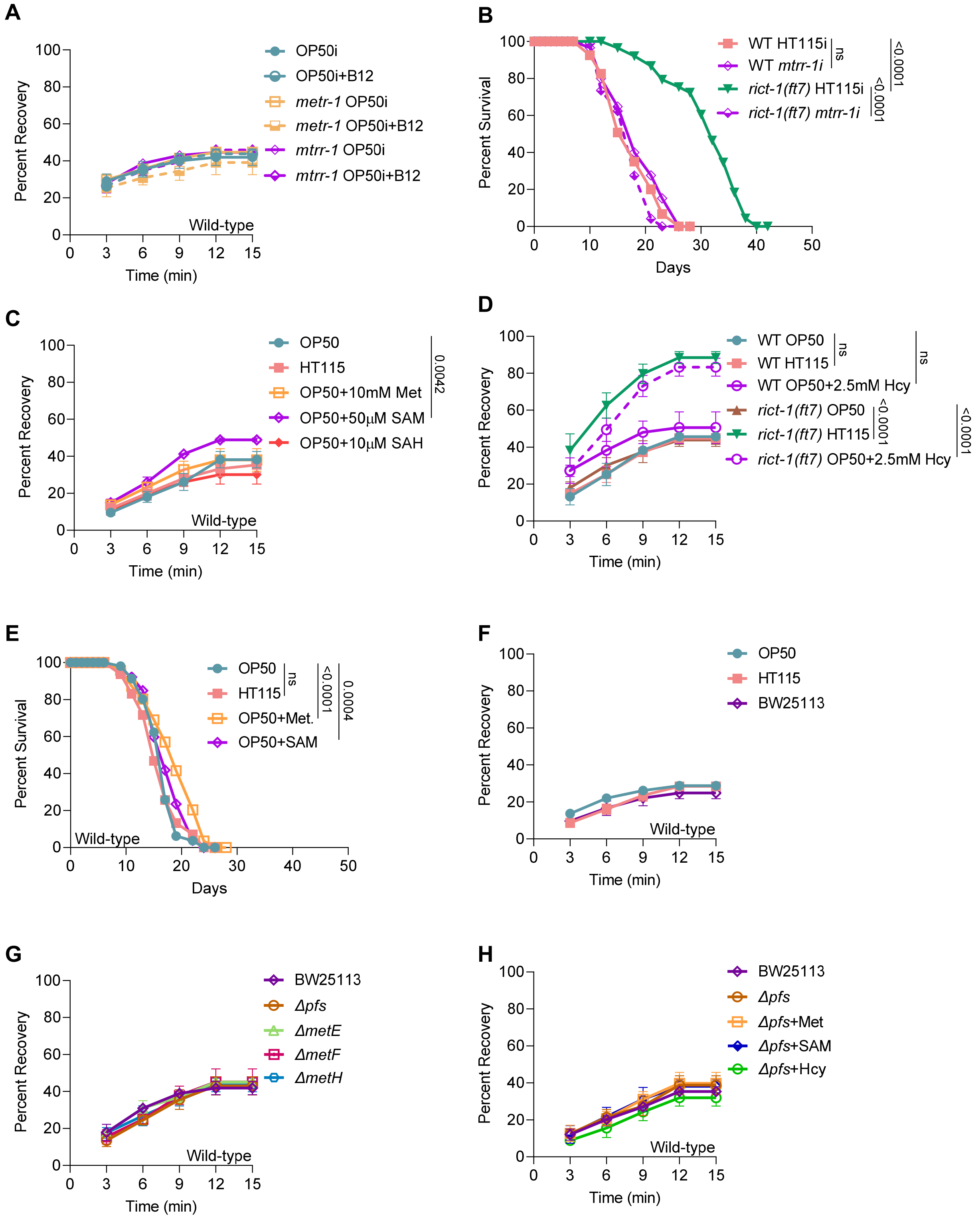


**Figure S2: (A)** Wild-type worms showed no effect on OST when *metr-1* or *mtrr-1* OP50i was supplemented with 64 nM B12. One of three biologically independent replicates is shown. P-value determined using Ordinary Two-way ANOVA with Tukey's multiple comparisons test and found to be non-significant. **(B)** A reduction in the lifespan of *rict-1(ft7)* worms was observed upon the knockdown of *mtrr-1* gene using HT115i. One of two biologically independent replicates is shown. P-value determined using Mann-Whitney U test. **(C)** No changes were observed in OST of wild-type worms upon supplementation of Met and SAH to the OP50 diet. Addition of SAM resulted in a small but significant increase in the OST of wild-type worms. One of two biologically independent replicates is shown. P-value determined using Ordinary Two-way ANOVA with Tukey's multiple comparisons test. **(D)** The *rict-1(ft7)* worms fed with Hcy-supplemented OP50 showed an increase in the OST while wild-type remained unaffected. One of two biologically independent replicates is shown. P-value determined using Ordinary Two-way ANOVA with Tukey's multiple comparisons test. **(E)** Addition of Met or SAM to the OP50 diet resulted in a small but significant increase in the lifespan of wild-type worms. One of two biologically independent replicates is shown. P-value determined using Mann-Whitney U test. **(F)** Wild-type worms showed no change in OST when grown on *E. coli* BW25113. One of three biologically independent replicates is shown. P-value determined using Ordinary Two-way ANOVA with Tukey's multiple comparisons test and found to be non-significant. **(G)** Wild-type worms exhibited no change in OST when grown on *E. coli* Δ*pfs,* Δ*metE*, Δ*metF*, or Δ*metH*. One of three biologically independent replicates is shown. P-value determined using Ordinary Two-way ANOVA with Tukey's multiple comparisons test and found to be non-significant. **(H)** Wild-type worms grown on Δ*pfs* exhibited no difference in OST on supplementation of Met, Hcy, or SAM. One of two biologically independent replicates is shown. P-value determined using Ordinary Two-way ANOVA with Tukey's multiple comparisons test and found to be non-significant. P-value ≥0.05 was considered not significant, ns. All experiments were performed at 20 °C. All data and analysis are provided in the Source Data file.

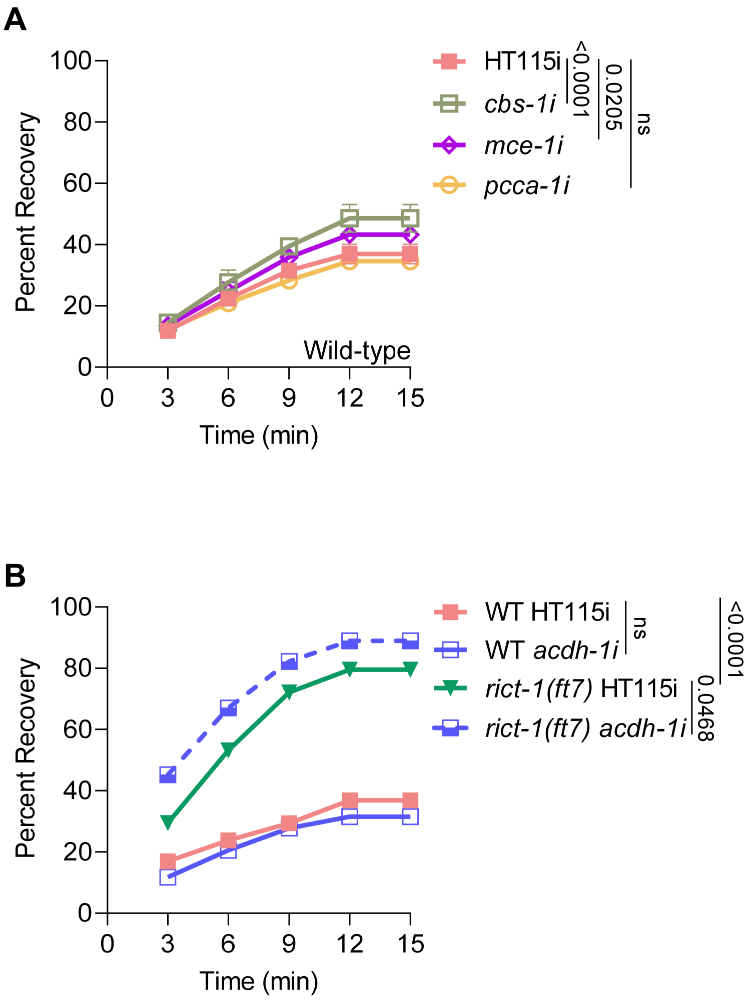


**Figure S3: (A)** Wild-type worms exhibited a small but significant change in OST upon the knockdown of *cbs-1* and *mce-1* using HT115i. One of three biologically independent replicates is shown. **(B)** The knockdown of *acdh-1* using HT115i did not affect the OST of *rict-1(ft7)* worms. One of three biologically independent replicates is shown. P-value determined using Ordinary Two-way ANOVA with Tukey's multiple comparisons test. P-value ≥0.05 was considered not significant, ns. All experiments were performed at 20 $^{o}$C. All data and analysis are provided in the Source Data file.

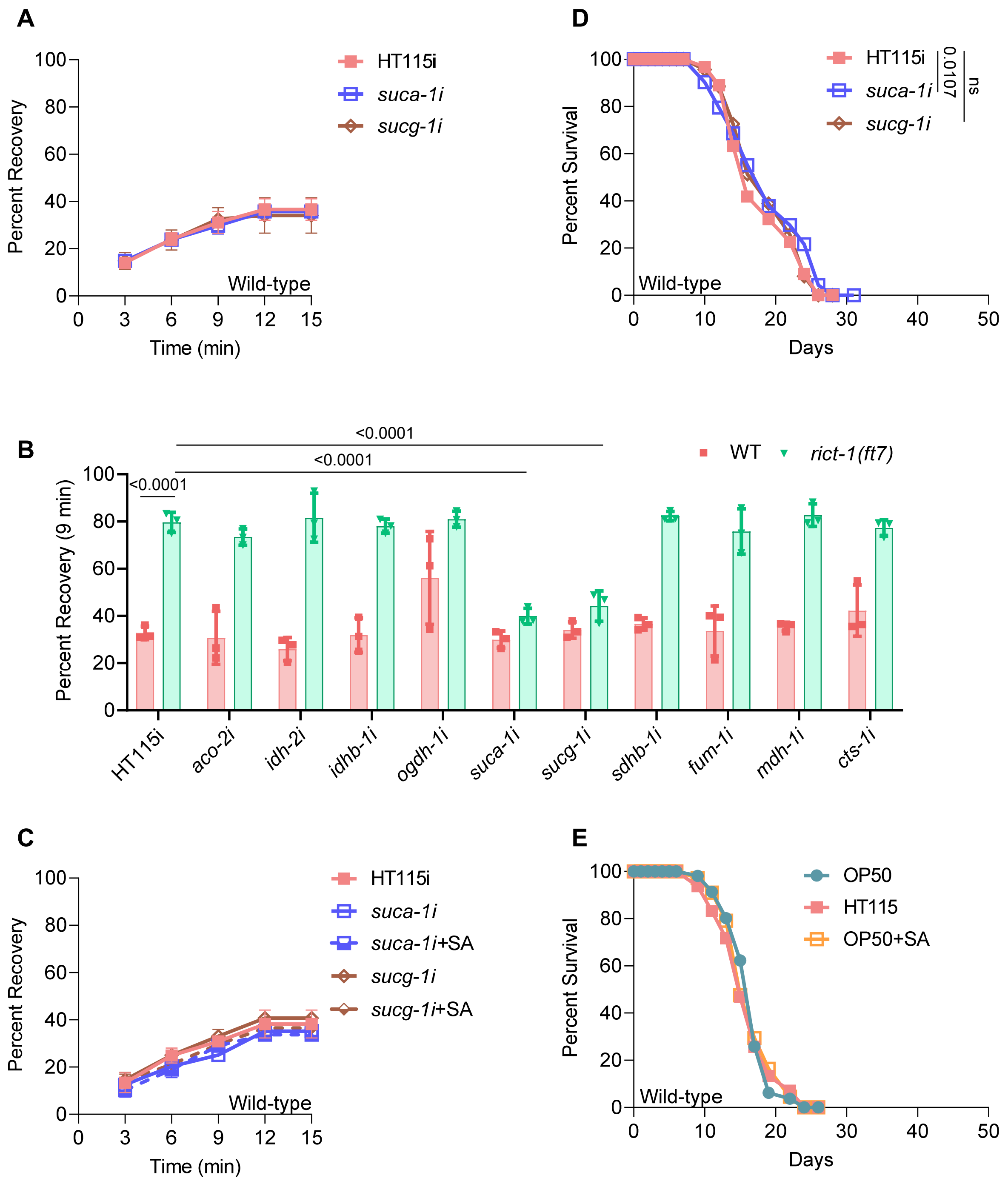


**Figure S4: (A)** Knockdown of *suca-1* or *sucg-1* using HT115i showed no effect on the OST of the wild-type worms. One of three biologically independent replicates is shown. P-value determined using Ordinary Two-way ANOVA with Tukey's multiple comparisons test and found to be non-significant. **(B)** Except for *suca-1* and *sucg-1*, knockdown of other genes of the TCA cycle did not affect the OST of the *rict-1(ft7)* worms. One of three biologically independent replicates is shown. P-value determined using Two-way ANOVA with Tukey's multiple comparisons test. **(C)** Supplementing SA to wild-type worms showed no change in the OST if *suca-1* or *sucg-1* was knocked down using HT115i. One of two biologically independent replicates is shown. P-value determined using Ordinary Two-way ANOVA with Tukey's multiple comparisons test and found to be non-significant. **(D)** Wild-type worms exhibited a small but significant change in the lifespan upon knockdown of *suca-1*, and no change was observed on knockdown of *sucg-1* using HT115i. One of three biologically independent replicates is shown. P-value determined using Mann-Whitney U test. **(E)** No difference in the lifespan was observed when wild-type worms were grown on OP50 supplemented with SA. One of two biologically independent replicates is shown. P-value determined using Mann-Whitney U test and found to be non-significant. P-value ≥0.05 was considered not significant, ns. All experiments were performed at 20 $^{o}$C. All data and analysis are provided in the Source Data file.

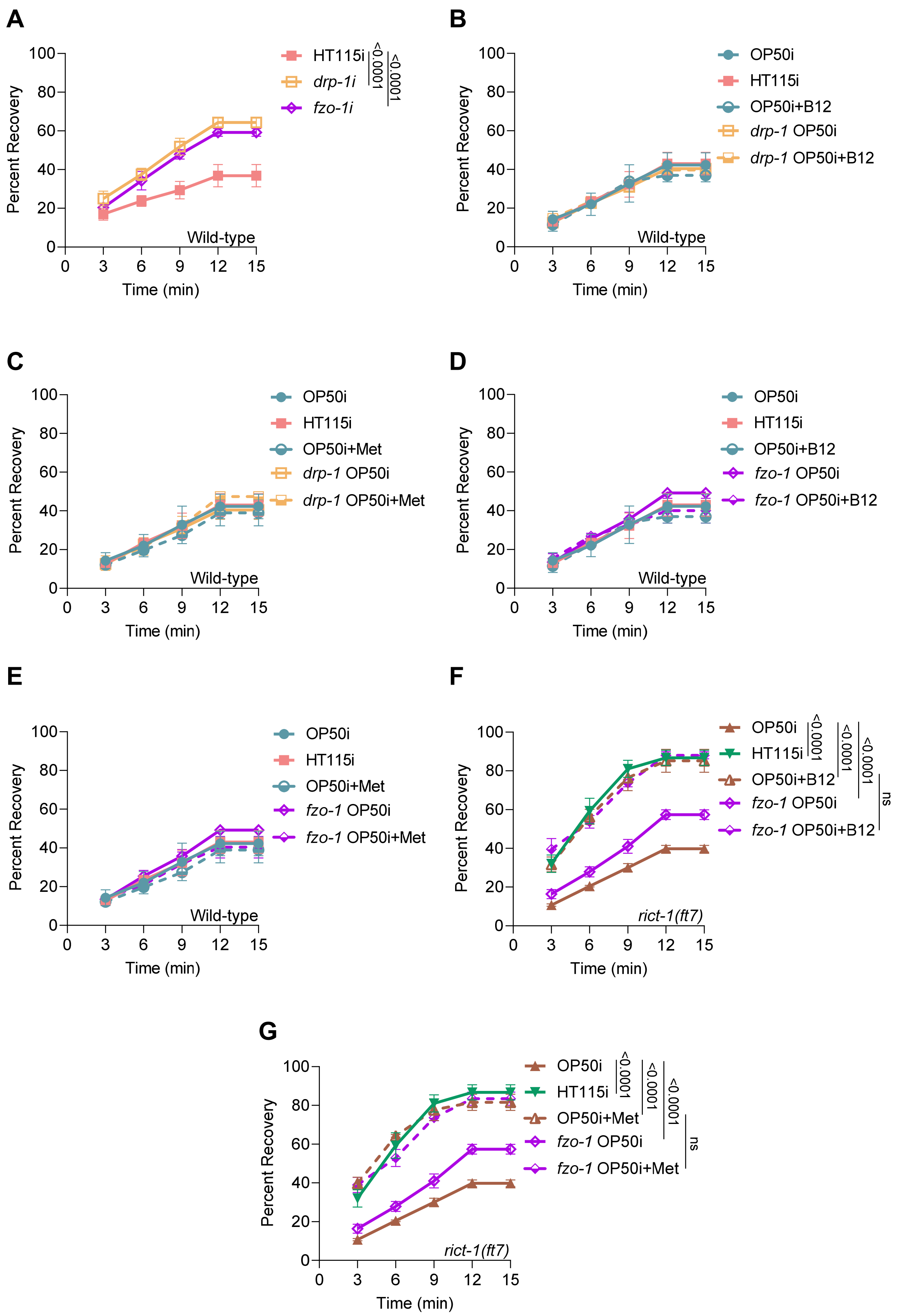
A
Percent Recovery
Time (min)
Wild-type
HT115i
drp-1i
fzo-1i
<0.0001
<0.0001
B
Percent Recovery
Time (min)
Wild-type
OP50i
HT115i
OP50i+B12
drp-1 OP50i
drp-1 OP50i+B12
C
Percent Recovery
Time (min)
Wild-type
OP50i
HT115i
OP50i+Met
drp-1 OP50i
drp-1 OP50i+Met
D
Percent Recovery
Time (min)
Wild-type
OP50i
HT115i
OP50i+B12
fzo-1 OP50i
fzo-1 OP50i+B12
E
Percent Recovery
Time (min)
Wild-type
OP50i
HT115i
OP50i+Met
fzo-1 OP50i
fzo-1 OP50i+Met
F
Percent Recovery
Time (min)
rict-1(ft7)
OP50i
HT115i
OP50i+B12
fzo-1 OP50i
fzo-1 OP50i+B12
<0.0001
<0.0001
<0.0001
ns
G
Percent Recovery
Time (min)
rict-1(ft7)
OP50i
HT115i
OP50i+Met
fzo-1 OP50i
fzo-1 OP50i+Met
<0.0001
<0.0001
<0.0001
ns

1

**Figure S5: (A)** Wild-type worms showed a modest increase in OST on knockdown of *drp-1* and *fzo-1* using HT115i. **(B)** No change in OST of wild-type worms was observed on knockdown of *drp-1* using OP50i supplemented with B12. **(C)** No change in OST of wild-type worms was observed on knockdown of *drp-1* using OP50i supplemented with Met. **(D)** No change in OST of wild-type worms was observed on knockdown of *fzo-1* using OP50i supplemented with B12. **(E)** No change in OST of wild-type worms was observed on knockdown of *fzo-1* using OP50i supplemented with Met. **(F)** Knockdown of *fzo-1* using OP50i supplemented with B12 showed no effect on the increased OST of *rict-1(ft7)*. **(G)** Knockdown of *fzo-1* using OP50i supplemented with Met showed no effect on the increased OST of *rict-1(ft7)*. One of three biologically independent replicates is shown for all the experiments. P-value determined using Ordinary Two-way ANOVA with Tukey's multiple comparisons test. P-value ≥0.05 was considered not significant, ns. All experiments were performed at 20 $^{o}$C. All data and analysis are provided in the Source Data file.

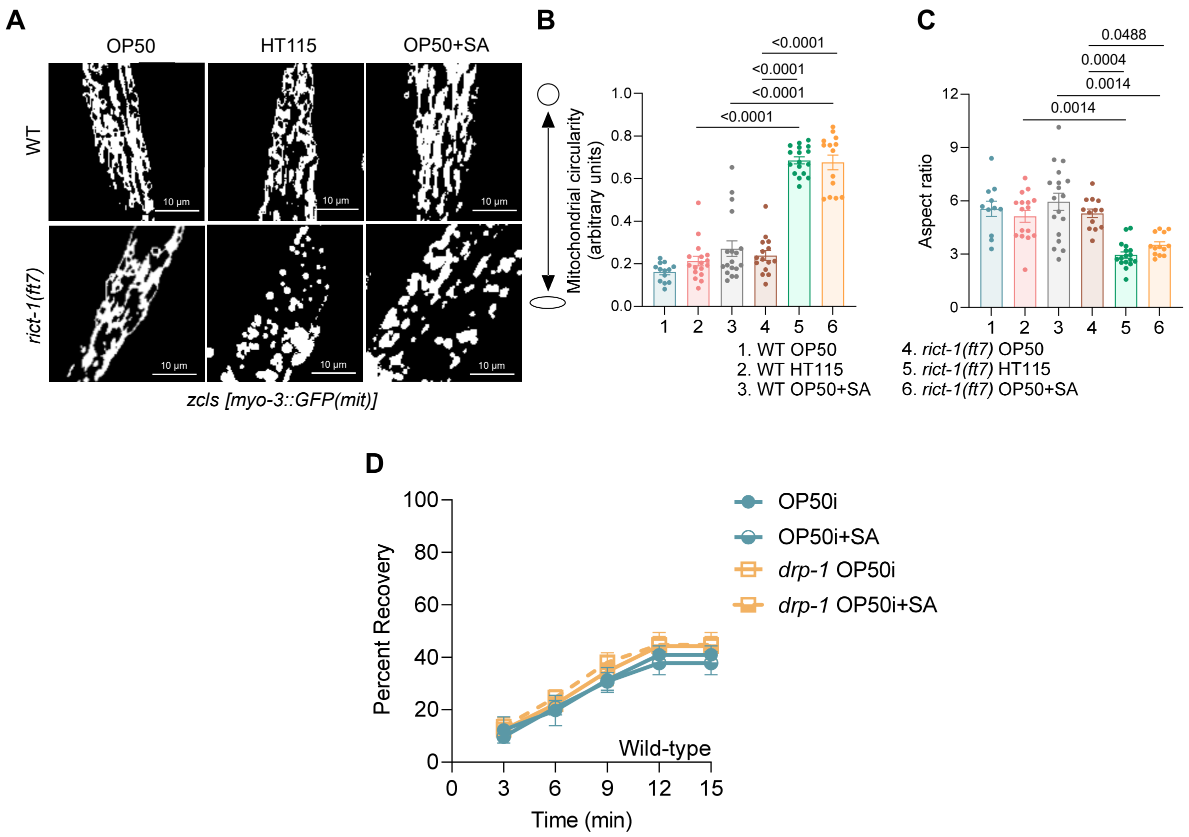


**Figure S6: (A-C)** Supplementation of SA to OP50 induced fragmentation of mitochondria in *rict-1(ft7)*, similar to that on the HT115 diet. (A) Representative images, (B) Mitochondrial circularity increases, and (C) aspect ratio reduces in the *rict-1(ft7)* worms grown on OP50 supplemented with SA. One of two biologically independent replicates is shown. P-value determined using Ordinary Two-way ANOVA with Tukey's multiple comparisons test. **(D)** The OST of wild-type worms remains unaffected when SA supplementation is combined with *drp-1i*. One of three biologically independent replicates is shown. P-value determined using Ordinary Two-way ANOVA with Tukey's multiple comparisons test and found to be non-significant. P-value ≥0.05 was considered not significant, ns. All experiments were performed at 20 °C. All data and analysis are provided in the Source Data file.

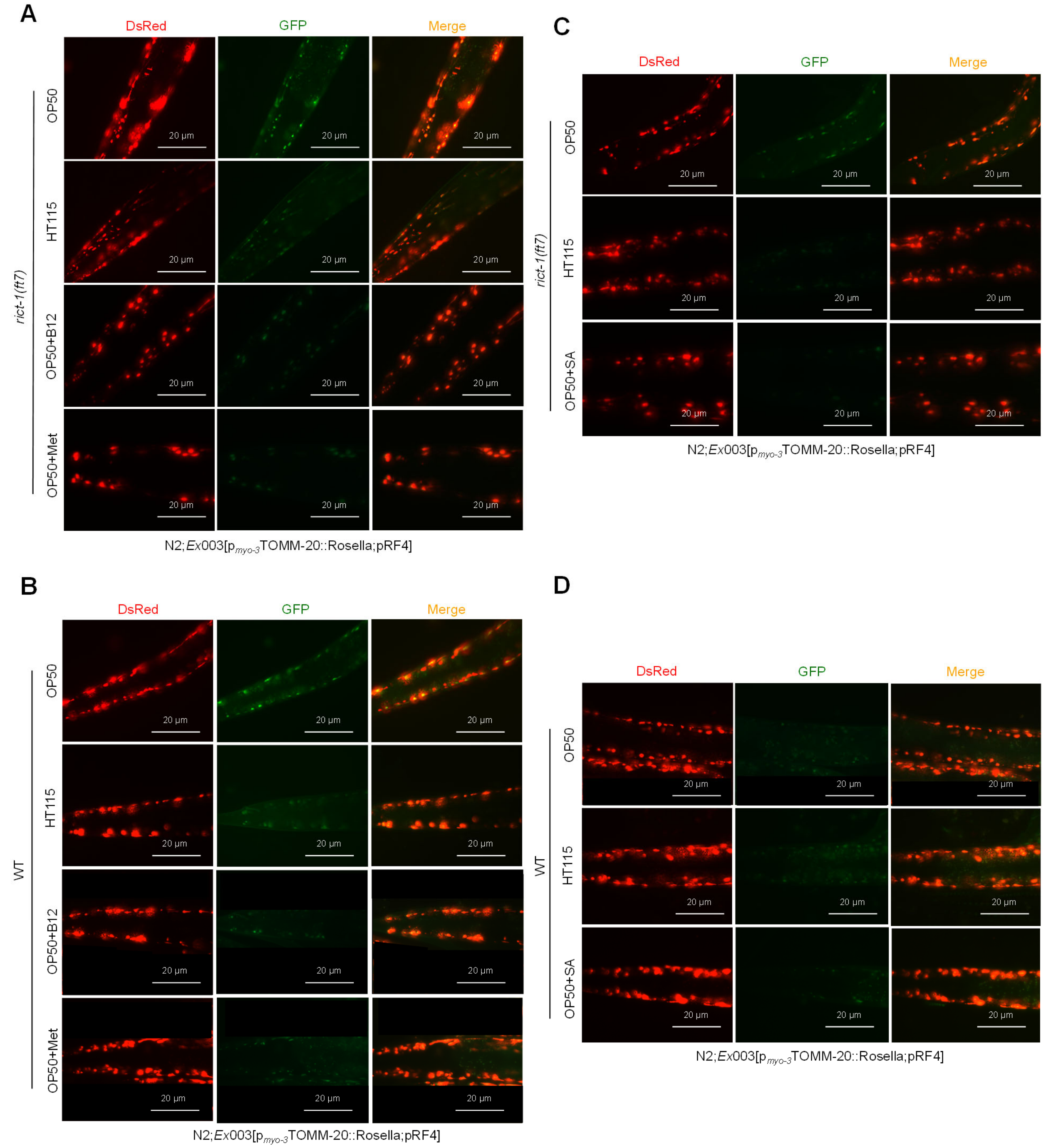


**Figure S7: (A and B)** Representative images of (A) *rict-1(ft7)* and (B) wild-type worms fed on B12 or Met-supplemented OP50 diet. **(C and D)** Representative images of (C) *rict-1(ft7)* and (D) wild-type worms fed on SA-supplemented OP50 diet. Images captured at 40X magnification. One of three biologically independent replicates is shown. All experiments were performed at 20 °C. Quantified data (presented in Figure 8B-D) and analysis for all conditions are provided in the Source Data file.

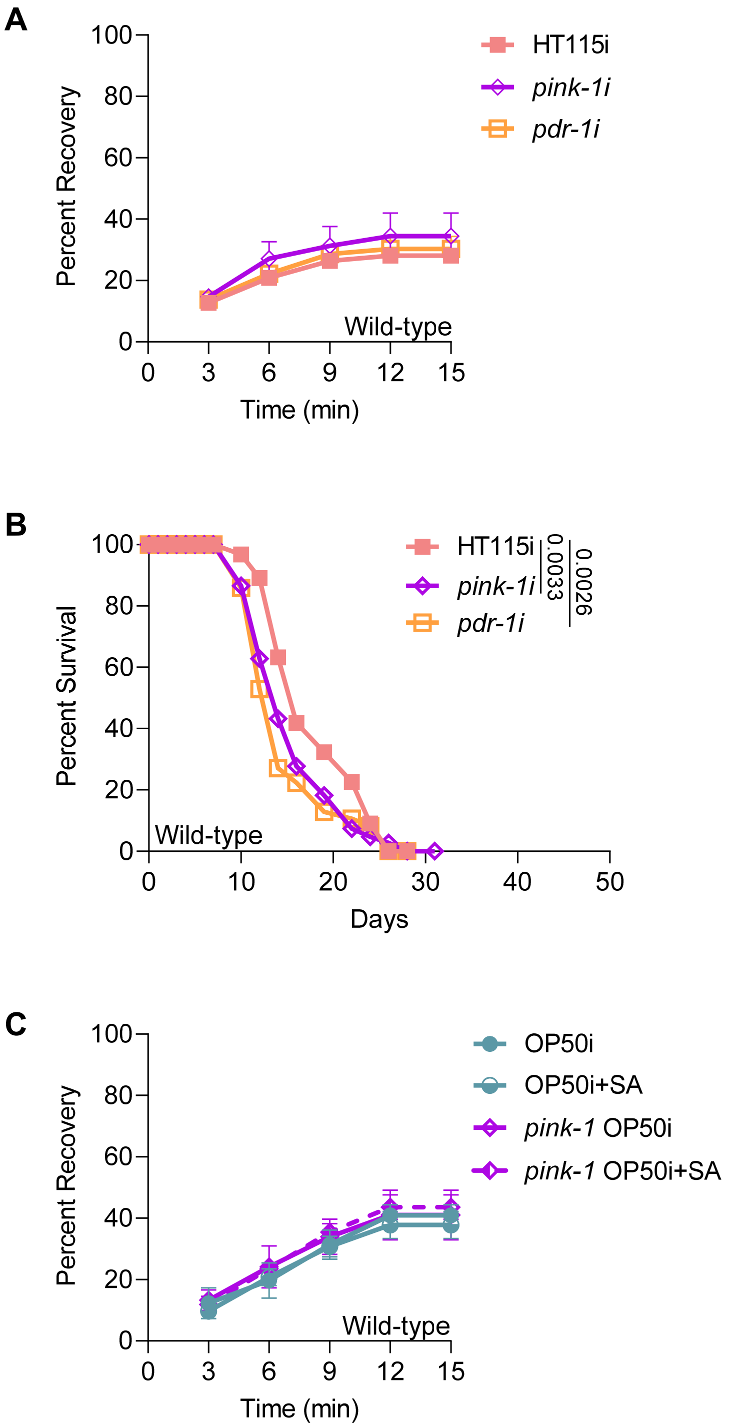


**Figure S8: (A)** Wild-type worms exhibited no change in OST on *pink-1* or *pdr-1* HT115i. One of three biologically independent replicates is shown. P-value determined using Ordinary Two-way ANOVA with Tukey's multiple comparisons test and found to be non-significant. **(B)** Wild-type worms exhibited a small reduction in lifespan upon knockdown of *pink-1* or *pdr-1 using* HT115i. One of three biologically independent replicates is shown. P-value determined using Mann-Whitney U test. **(C)** Wild-type worms showed no effect on supplementation of SA in *pink-1* OP50i. One of three biologically independent replicates is shown. P-value determined using Ordinary Two-way ANOVA with Tukey's multiple comparisons test and found to be non-significant. All experiments were performed at 20 $^{o}$C. All data and analysis are provided in the Source Data file.
